# MHC-I downregulation is associated with immune-evasion in breast cancers with heterogeneous activation of epithelial-mesenchymal plasticity

**DOI:** 10.64898/2026.10.17.759751

**Authors:** Kimaya Bakhle, Andreas Stephanou, Sarthak Sahoo, Ioannis Ntekas, Kaden Priebe, Mayisa Rahman, Rishi Puri, Scott Butler, Peter Schweitzer, Shiney Chandraganti, Mohit K. Jolly, Iwijn De Vlaminck, Praveen Sethupathy, Anushka Dongre

## Abstract

Cancer cell-intrinsic epithelial-mesenchymal plasticity (EMP) drives carcinoma progression through increased intratumoral heterogeneity and an immunosuppressive tumor microenvironment (TME). Specifically, EMP activation in murine mammary carcinomas results in exclusion of CD8^+^ T-cells and recruitment of immunosuppressive M2-like macrophages, resulting in resistance to anti-CTLA-4 immune checkpoint blockade (ICB) therapy. Notably, mixed E-qM tumors in which only 10% of cancer cells have activated EMP are resistant anti-CTLA-4 immune checkpoint blockade (ICB) therapy. This suggests that minority fractions of qM cancer cells could, in theory, exert immunosuppressive effects on surrounding E cancer and immune cells to shield the entire heterogeneous tumor against immune attack. However, this has not yet been demonstrated. Here, we use spatial transcriptomic profiling to demonstrate that mixed E-qM tumors assemble immunosuppressive cellular niches. Furthermore, E cancer cells within mixed E-qM tumors experience long-term MHC-I downregulation leading to resistance to elimination by CD8^+^ T-cells. This is associated with the exclusion of T-cells from the tumor core and a dysfunctional phenotype in the remaining CD8^+^ T-cells. qM cancer cells recruit immunosuppressive M2-like macrophages to mixed tumors, which further downregulates E cancer cell-intrinsic MHC-I levels. Finally, we demonstrate that human triple-negative breast cancers with increased EMP activation exclude T-cells from the tumor core. Together, these findings reveal that qM cancer cells modulate the immune infiltrate of mixed tumors, promoting MHC-I downregulation and evasion of immunological control by CD8^+^ T-cells.

**Significance statement:** Epithelial-mesenchymal plasticity (EMP) drives the formation of an immunosuppressive tumor microenvironment (TME). Notably, EMP activation in just 10% of cancer cells promotes immunosuppression and immunotherapy resistance in heterogeneous murine mammary tumors. However, it was previously unknown how minority fractions of EMP-activated cancer cells act within heterogeneous tumors to cross-protect neighboring cancer cells against immune attack. Here, we show that minority fractions of quasi-mesenchymal cancer cells result in an altered immune infiltrate with fewer T-cells and increased tumor-associated macrophages. Together, these changes to the TME lead to MHC-I downregulation in surrounding epithelial cancer cells, which shields these cells against recognition by cytotoxic CD8^+^ T-cells. These results reveal a potential mechanism of immune-evasion and immunotherapy resistance in heterogeneous breast tumors.

## Introduction

Breast cancer is a heterogeneous disease, both at the population level (intertumoral heterogeneity) and the single patient level (intratumoral heterogeneity)^1–8^. One driver of intratumoral heterogeneity is the diverse tumor microenvironment (TME). This dynamic collection of cancer, immune, and stromal cells can regulate disease progression and therapy response^9,10^. For example, pro-tumor cells (e.g. Tregs, M2-like macrophages) can blunt the cytolytic effects of anti-tumor immune cells (e.g. CD8^+^ T-cells, NK cells, M1-like macrophages), rendering them incapable of eliminating tumors^11–13^. Patients with fewer tumor-infiltrating anti-tumor immune cells have poor prognoses and fail to respond to therapeutics, such as immune checkpoint blockade therapy^14–20^.

Cancer-cell intrinsic causes of intratumoral heterogeneity include genomic instability, neoantigen heterogeneity, epigenetic regulation, metabolic adaptation, treatment resistance, and cellular plasticity^1,9,21,22^. One type of cellular plasticity that contributes to intratumoral heterogeneity is epithelial-mesenchymal plasticity (EMP)^23–26^. EMP is a cellular program that enables epithelial cells, which express the markers EpCAM and E-cadherin, to gain mesenchymal markers, such as vimentin, and N-cadherin, and features including invasiveness, metastatic potential, and stem-cell like properties^27–37^. These changes are driven by EMP-inducing transcription factors (EMP-TFs) Zeb1, Twist, and Snail. EMP induction results in a spectrum of hybrid or quasi-mesenchymal (qM) plastic states, which share phenotypic characteristics and markers of both epithelial and mesenchymal cells.^37,38^. These hybrid states display enhanced tumorigenicity^39–41^. Moreover, this is a reversible process, as hybrid or qM cells can undergo a mesenchymal-epithelial transition and return to an epithelial-like state^37^.

We and others have demonstrated that EMP is a driver of immunosuppression in the TME of breast, hepatocellular, and pulmonary carcinomas, as well as melanoma^42–48^. Specifically, we used MMTV-PyMT-Snail-YFP mice to isolate EpCAM^hi^ Snail-YFP^lo^ epithelial (E) and EpCAM^lo^ Snail-YFP^hi^ qM cell lines^43^. We then reimplanted E and qM cells into separate syngeneic hosts to generate completely E or qM tumors. We found that E tumors recruited immunopermissive CD8^+^ T-cells to the tumor center and restricted M1-like macrophages to the tumor periphery. Conversely, qM mammary tumors excluded T-cells to the periphery and instead recruited high numbers of M2-like macrophages and Tregs to the tumor core. These differences in the TME of E and qM tumors also led to differences in response to anti-CTLA-4 immune checkpoint blockade (ICB) therapy, where epithelial tumors were sensitive to this therapy and quasi-mesenchymal tumors were resistant^43^. These findings demonstrate that EMP-mediated immunosuppression is a driver of immunotherapy resistance in breast cancers, among other solid tumors.

However, these findings come from tumors consisting of homogeneous cell populations, which is not representative of the heterogeneity observed in breast cancer patient samples^1,2,8,49–51^. To better recapitulate this heterogeneity, we generated mixed E-qM tumors containing a minority fraction (10%) of qM cells. Notably, these mixed tumors were resistant to anti-CTLA-4 ICB therapy. Upon microscopic evaluation of these tumors, we found that the E and qM cancer cells segregate to distinct regions within the mixed tumor^43^. This topographical separation suggests that qM cancer cells release paracrine factors that act on E and immune cells within mixed tumors to protect the entire tumor against immune attack. This phenomenon, which we term cross-protection, is clinically relevant, as human breast cancers contain minority fractions of qM cancer cells that have the potential to drive resistance of the entire tumor to ICB therapy^52–56^. However, it is unknown how factors released by qM cancer cells alter surrounding epithelial and immune cells to drive cross-protection against immune attack.

We hypothesized that qM cancer cells cross-protect neighboring E cancer cells either through direct changes or indirectly through alterations to immune cells within the TME of mixed tumors. To understand cross-protection, we first investigated the spatial range of cross-protection. We found that separating qM cancer cells into a contralateral tumor reversed cross-protective effects, suggesting that local paracrine signaling within the TME is required. Next, we investigated immunological mechanisms of E tumor control. We found that CD8^+^ T-cells constrain E tumor growth, but this ability is lost in mixed tumors containing just 10% qM cancer cells. The inability of CD8^+^ T-cells to control mixed tumor growth could be attributed to a combination of T-cell exclusion and decreased CD8^+^ T-cell functionality. The altered location and function of CD8^+^ T-cells also led to decreased E cancer cell-intrinsic major histocompatibility complex (MHC)-I expression within mixed tumors. We found that MHC-I expression on E cancer cells was also decreased in mixed tumors in the absence of lymphocytes. Therefore, we investigated macrophages as a regulator of MHC-I expression. We found that depletion of macrophages produced a trend towards restored MHC-I expression on E cancer cells. Finally, we identified that human triple-negative breast cancers (TNBC) with increased EMP activation display T-cell exclusion. These findings suggest that within mixed tumors, qM cancer cells generate an immunosuppressive TME, which prevents MHC-I induction on E cancer cells, leading to immune evasion and ICB resistance.

## Results

### Cross-protection occurs locally rather than systemically

We have previously observed that mixed pB2:Snail^hi^ (9:1) tumors are partially resistant to anti-CTLA-4 therapy^43^. Additionally, we have observed that untreated pB2:Snail^hi^ tumors grow significantly faster when implanted in syngeneic hosts compared to pB2 tumors **(Fig S1A)**. Therefore, minority fractions of qM cancer cells likely confer pro-tumorigenic and immunosuppressive attributes to E cancer cells within mixed tumors.

We first sought to understand whether the physical juxtaposition of epithelial and quasi-mesenchymal cancer cells within a given primary tumor is required for cross-protection against immune attack. Accordingly, we separated the two cell types into contralateral tumors located at distinct anatomical sites. To achieve this, we used previously-established E and qM cell lines derived from mouse mammary tumor virus-polyoma middle T antigen (MMTV-PyMT) tumor-bearing mice containing a Snail-YFP fluorescent reporter^43^. Briefly, late-stage tumors were excised from female MMTV-PyMT mice, and the tumor cells were sorted based on the epithelial marker EpCAM and expression of the Snail-YFP reporter. This resulted in the E cell lines pB2 and EpCAM^hi^ and the qM cell line Snail^hi^ ^35,42,43,57^.

We implanted pB2 cells in the right mammary fat pad of a female C57BL/6J mouse and either Matrigel (Mg) alone or containing Snail^hi^ cells into the left mammary fat pad of the mouse **(Fig 1A)**. We allowed the tumors to grow until the Snail^hi^ tumors reached 2000 mm^3^ and then collected the pB2 tumors to assess changes to the TME in the presence of a contralateral qM tumor. By flow cytometry, we found that the percentage of CD8^+^ T-cells and their expression of the activation markers IFN-γ, TNF-α, Perforin, and Granzyme B were unaffected by the presence of a contralateral Snail^hi^ tumor **(Fig 1B, S1B-C)**. CD8^+^ T-cells under both conditions maintained low expression of the dysfunction markers TIM3, TIGIT, PD-1, and CTLA-4 **(Fig 1C, S1B-C)**. Additionally, the percentage of CD4^+^ T-cells, regulatory T-cells (Tregs), macrophages, and monocytes showed no difference in pB2 tumors contralateral to either Mg or a qM tumor **(Fig 1D-F, S1D-E)**. pB2 tumors contralateral to Snail^hi^ tumors did show an increase in granulocytes **(Fig 1F).** To confirm that implanted E tumors do not activate EMP or become seeded by contralateral qM cells^58^, we stained pB2 tumors for the epithelial marker E-cadherin and mesenchymal marker vimentin. We found that pB2 tumors remained epithelial under either condition, with high expression of E-cadherin and low expression of vimentin **(Fig 1G)**. We found that CD3^+^ T-cells infiltrated the pB2 tumors under either condition and macrophages remained restricted to the tumor periphery **(Fig 1H, 1J).** The macrophages expressed low levels of Arginase-1 (Arg1), likely indicating a M1-like polarization state **(Fig 1I-J).** Finally, the pB2 tumors remained responsive to anti-CTLA-4 therapy with the presence of a contralateral Snail^hi^ tumor **(Fig 1K-L, S1F)**. Together, these findings demonstrate that physical proximity of E and qM cancer cells is required for cross-protection. Upon separation of qM cancer cells, E tumors maintain an immunopermissive TME and sensitivity to anti-CTLA-4 therapy.

**Figure 1.**
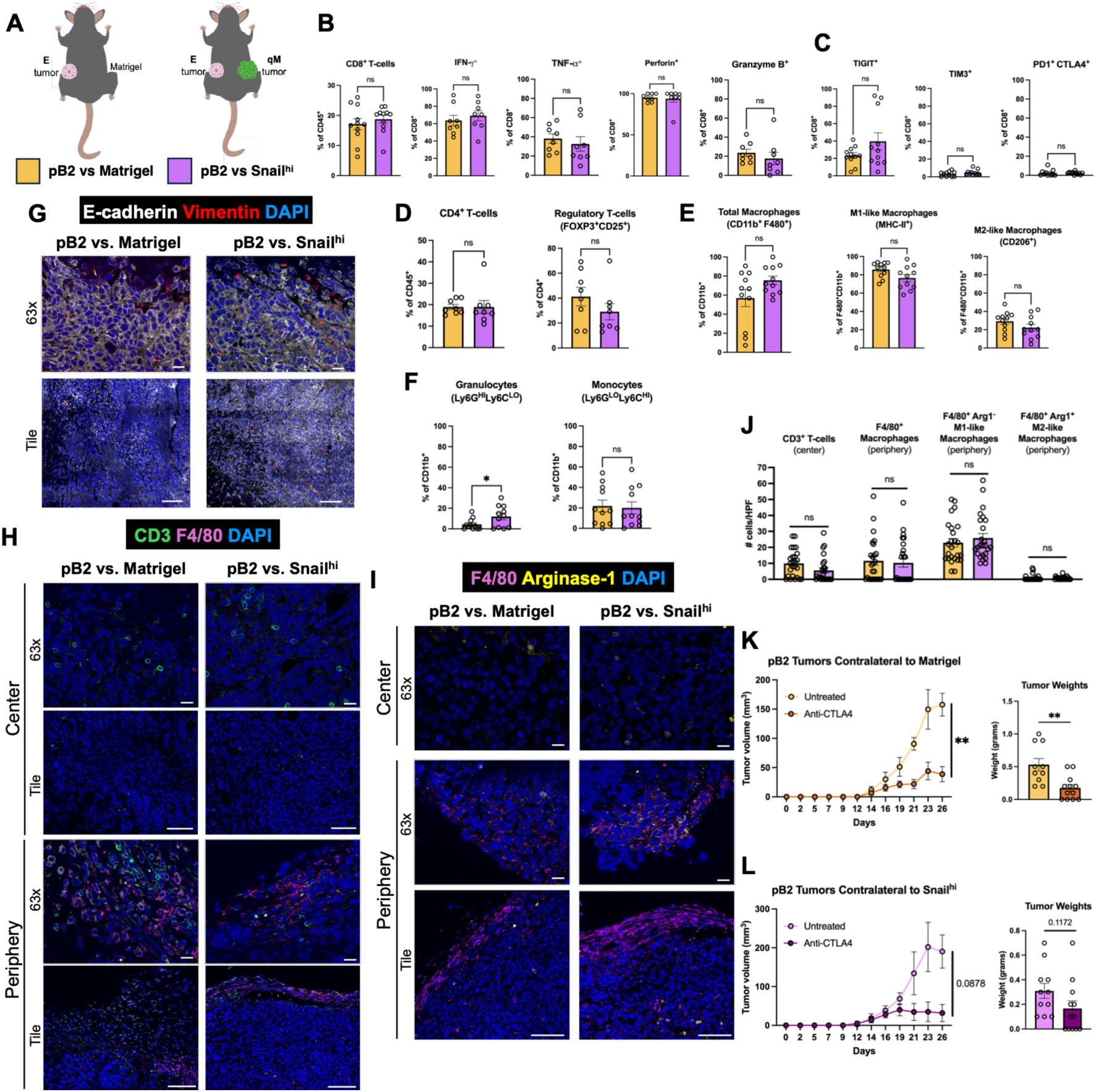
Cross-protection is lost upon physical separation of epithelial and quasi-mesenchymal cancer cells. **(A)** Epithelial pB2 cells were implanted contralateral to matrigel (Mg, yellow) or Snail^hi^ cells (purple). **(B)** Percentage of CD8^+^ T-cells expressing activation markers, as measured by flow cytometry. **(C)** Percentage of CD8^+^ T-cells expressing dysfunction markers. **(D)** Percentage of CD4^+^ T-cells and regulatory T-cells. **(E)** Percentage of macrophage subpopulations. **(F)** Percentage of granulocytes and monocytes. **(G)** Immunofluorescent staining of EMP markers E-cadherin (white) and Vimentin (red), along with DAPI (blue), demonstrating that pB2 tumors remain epithelial when implanted contralateral to a Snail^hi^ tumor. Scale bars indicate 20 μm in 63x images and 100 μm in tile images. **(H)** Immunofluorescent staining showing that CD3^+^ T-cells (green) infiltrate pB2 tumors and F4/80^+^ macrophages (magenta) are restricted to the periphery under both conditions. **(I)** F4/80^+^ Macrophages (magenta) at the periphery of pB2 tumors contralateral to Snail^hi^ tumors retain low arginase-1 (yellow) expression, indicating an M1-like state. Scale bars indicate 20 μm in 63x images and 100 μm in tile images in H-J. **(J)** Quantification of immune cells across five high power fields in five tumors for each condition. **(K)** Tumor kinetics of pB2 tumors implanted contralateral to Mg and left untreated or treated with anti-CTLA-4. **(L)** Tumor kinetics of pB2 tumors implanted contralateral to Snail^hi^ tumors. **B-F** represent three independent experimental replicates including *n*=3-4 per group. Data are represented as mean ± standard error of the mean (SEM) and were analyzed using an unpaired two-tailed t-test. **G-J** represent two independent experimental replicates including *n*=5 total per group. Data are represented as mean ± standard error of the mean (SEM) and were analyzed using an unpaired two-tailed t-test. **K-L** represent three independent experimental replicates including *n*=3-4 per group. Data are represented as mean ± standard error of the mean (SEM) and were analyzed using a mixed-effects model (for tumor kinetics) and unpaired two-tailed t-test (for tumor weights). In **B-F** and **J-L**, \**p* < 0.05 and \*\**p* < 0.01. Comparisons are not significant (ns, *p* ≥ 0.05) if not indicated.

To determine whether these findings could be generalized to other models, we repeated this experimental setup using a second E cancer cell line: EpCAM^hi^ **(Fig S2A)**. We found that CD8^+^ T-cells remained functional with high expression of effector molecules and low expression of dysfunction markers in EpCAM^hi^ tumors contralateral to Mg or Snail^hi^ tumors (**Fig S2B-D)**. The percentage of Tregs and myeloid cells were also unchanged under both conditions (**Fig S2E-H).** By immunofluorescent staining, we confirmed that the EpCAM^hi^ tumor cells remained epithelial upon implantation. There were few tumor-infiltrating CD3^+^ T-cells in EpCAM^hi^ tumors contralateral to either Mg or Snail^hi^ tumors, but the number of F4/80^+^ Arg1^-^ M1-like and F4/80^+^ Arg1^+^ M2-like macrophages were similar under both conditions **(S2I)**. Although not to the same extent as pB2 tumors, EpCAM^hi^ tumors were responsive to anti-CTLA-4 when implanted contralateral to either Mg or a Snail^hi^ tumor (**Fig S2J)**.

Finally, we profiled circulating T-cells and myeloid cells to determine if qM-derived factors can induce systemic immunosuppression. We found that circulating CD8^+^ T-cells show low expression of the dysfunction markers TIM3, TIGIT, PD-1, and CTLA-4 in the blood and tumor-draining lymph-node of mice bearing pB2 tumors contralateral to Mg or Snail^hi^ tumors (**Fig S3A-D)**, likely indicating retained effector function. Percentages of myeloid cells were also unaffected systemically with implantation of either Mg or a Snail^hi^ tumor **(Fig S3E-F).** These findings were consistent in circulating immune cells in mice bearing EpCAM^hi^ tumors contralateral to Mg or Snail^hi^ tumors **(Fig S3G-L)**.

Taken together, our findings from pB2 and EpCAM^hi^ E cancer cells implanted contralateral to Snail^hi^ qM cancer cells demonstrate that cross-protection requires physical proximity, specifically within the same tumor. pB2 and EpCAM^hi^ cells separated from qM cancer cells remain immunopermissive and responsive to treatment with anti-CTLA-4. Additionally, circulating immune cells express low levels of immunosuppressive markers, suggesting that qM tumors do not induce systemic immunosuppression. Therefore, we concluded that local, rather than systemic, paracrine signaling is required for qM cancer cells to cross-protect E cells against immune attack.

### Minority fractions of qM cancer cells confer E cancer cells with resistance to elimination by CD8^+^ T-cells

Given that cross-protection occurs within the confines of the primary tumor, we focused on mechanisms of immunosuppression within the local TME. We first sought to understand how E tumor growth is controlled, in order to understand how this is altered in mixed tumors. To determine whether lymphocytes control E tumor growth, we implanted pB2 tumors in immunocompetent C57BL/6J mice and immunodeficient *Rag1*^-/-^ mice. We found that pB2 tumors grew significantly faster and formed larger tumors in *Rag1*^-/-^ mice. Since pB2 cells express high levels of MHC-I^43^, we next wanted to determine if this effect could be attributed to CD8^+^ T-cells. Indeed, pB2 tumors implanted in *Cd8a*^tm1Mak^ (CD8-KO) mice formed significantly larger tumors than those implanted in C57BL/6J mice **(Fig 2A, S4A)**. There was no appreciable difference in growth of pB2 tumors between *Rag1*^-/-^ and CD8-KO mice, suggesting that CD8^+^ T-cells are the primary adaptive immune subset responsible for controlling the growth of E tumors. To determine whether CD8^+^ T-cells also control growth of mixed tumors, we repeated this experimental setup with mixed pB2:Snail^hi^ (9:1) tumors. Conversely, we found no difference in mixed tumor growth between the three hosts, suggesting that mixed tumors evade control by CD8^+^ T-cells **(Fig 2B, S4A)**. qM cancer cells remained the minority population in mixed tumors in both C57BL/6J and *Rag1*^-/-^ mice **(Fig S4B-D).**

**Figure 2.**
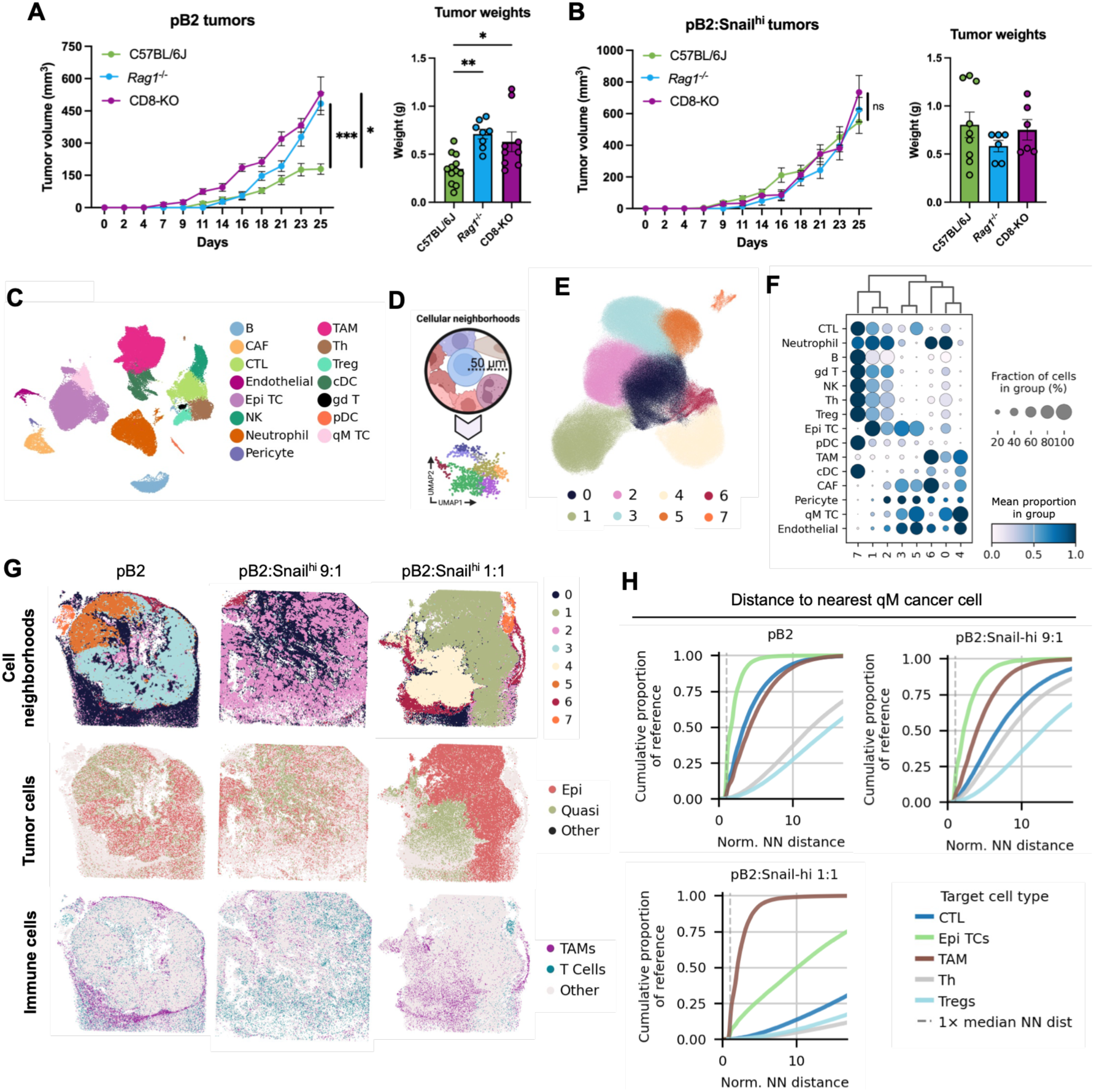
Mixed tumors evade control by CD8^+^ T-cells and form immunosuppressive niches. **(A-B)** Tumor kinetics and weights of epithelial and mixed tumors implanted in immunocompetent (C57BL/6J) and immunodeficient (*Rag1*^-/-^ and CD8-KO) mice. **(C)** Uniform manifold approximation and projection (UMAP) plot of merged public and in-house scRNA-seq data colored by cell type. **(D)** Overview of the neighborhood analysis, in which the local cell type composition of each cell was computed within a 50-μm radius, followed by dimensionality reduction and Leiden clustering. **(E)** Cellular neighborhoods presented in UMAP space. **(F)** Cell type composition within each cellular neighborhood. Color scale depicts the scaled mean proportion across neighborhoods for each cell type. Dot size represents the fraction of cells that have a non-zero presence of a given cell type in their local neighborhood. **(G)** Spatial distribution of cellular neighborhoods, cancer cells, and immune cells in epithelial and mixed tumors. **(H)** Normalized nearest-neighbor distances of select immune cell types to nearest qM cancer cell in E and mixed tumors. **A-B** represent two (*Rag1*^-/-^) or three (C57BL/6J, CD8-KO) independent experimental replicates including *n*=3-5 per group. Data are represented as mean ± standard error of the mean (SEM) and were analyzed using a mixed-effects model (for tumor kinetics) and one-way ANOVA (for tumor weights). \**p* < 0.05, \*\**p* < 0.01, \*\*\**p* < 0.001. Comparisons are not significant (ns, *p* ≥ 0.05) if not indicated. **C** includes 20 scRNA-seq datasets. **D-H** represent one experimental replicate per condition.

We next wanted to understand whether the location and function of immune cells are altered in mixed tumors. To this end, we performed single-cell RNA sequencing (scRNA-seq) and spatial transcriptomics of pB2 and pB2:Snail^hi^ (9:1 and 1:1) tumors. To supplement our newly generated scRNA-seq data, we incorporated publicly available mouse mammary tumor scRNA-seq datasets from scBaseCount, an Arc Institute resource in which public sequencing data are uniformly reprocessed from raw reads (^59^; see Methods). By scRNA-seq, we identified epithelial (Epi) and qM cancer cell populations, cancer-associated fibroblasts (CAFs), endothelial cells, pericytes, and several immune cell clusters including B cells, cytotoxic lymphocytes (CTLs; CD8^+^ T-cells), NK cells, neutrophils, tumor-associated macrophages (TAMs), T helper cells (Th; CD4^+^ T-cells), Tregs, gamma delta T-cells (gd T), conventional dendritic cells (cDCs), and plasmacytoid dendritic cells (pDCs) **(Fig 2C, S5A-D).**

Using the spatial transcriptomics data, we aimed to assess cancer-immune cell colocalization in an unbiased manner. To achieve this, we generated seven cellular neighborhoods **(Fig 2D-F, S5E)**. Epithelial tumor cells (Epi TCs) were primarily represented in clusters 1, 2, 3, and 5. Of these, clusters 1 and 2 demonstrated that E cancer cells co-localize with several immune cell types including neutrophils, B-cells, NK cells, Tregs, gd T-cells, cytotoxic lymphocytes (CTLs), and T helper (Th) cells. Quasi-mesenchymal cancer cells (qM TCs) were represented in clusters 0, 3, 4, and 5. Cluster 4 was depleted of immune cells, apart from TAMs and cDCs **(Fig 2F)**. The pB2 tumor was primarily composed of clusters 0, 2, 3, and 5. The pB2:Snail^hi^ 9:1 tumor contained clusters 0 and 2. Cells in the 1:1 mixed tumor primarily belonged to clusters 1, 4, and 6. Clusters 1 and 2 overlapped with E cancer cells and were enriched for lymphocytes. Clusters 0 and 4 overlapped with qM cancer cells and were enriched for TAMs and depleted of other immune cells **(Fig 2F-G)**. Using the Smoothie analysis pipeline^60^, we found that qM cancer cells overlap with a gene module representing TAM-related genes. Additionally, this region showed overlapping expression of CSF1R and its cognate ligands CSF1 and IL-34 **(Fig S5F)**. Finally, to empirically assess co-localization of immune cells with cancer cells, we calculated the normalized nearest neighbor distance of multiple immune cell types to qM cancer cells. We found that in all tumors, TAMs showed reduced distance from qM cancer cells, whereas CD4^+^ and CD8^+^ T-cells, as well as Tregs showed greater distances. As we observed visually, there was greater separation of E and qM cancer cells in the pB2:Snail^hi^ 1:1 sample compared to the pB2 and pB2:Snail^hi^ 9:1 samples **(Fig 2H).** Based on these results, we concluded that E and qM cancer cells create distinct immune cell niches within mixed tumors.

### Epithelial cancer cells within mixed tumors downregulate surface expression of MHC-I

After observing that CD8^+^ T-cells can control the growth of epithelial tumors, but not mixed tumors, we wanted to ask how E cancer cells are altered within mixed tumors to promote immune-evasion. Since CD8^+^ T-cells recognize target cells through peptide presented on MHC-I, we measured surface H-2Kb expression. By flow cytometry, we found that CD45^neg^ EpCAM^hi^ Snail-YFP^neg^ E cancer cells **(Fig S6A)** within mixed tumors had reduced H-2Kb expression compared to cancer cells from epithelial tumors **(Fig 3A, S6B)**. Using the spatial transcriptomics data of pB2 and pB2:Snail^hi^ (9:1 and 1:1) mixed tumors, we visualized the spatial distribution of MHC-I expression in cancer cells. As expected, *H2-K1* expression (encoding MHC-I molecule H-2Kb) was more pronounced in E cancer cells compared to qM cancer cells within the same mixed tumor^43^ **(Fig 3B)**. In the pB2:Snail^hi^ 1:1 tumor, expression of several MHC-I genes (*H2-K1, H2-D1, H2-Q7)* was decreased in E cancer cells located proximal to the qM region. Additionally, regulators of MHC-I expression and function including *Tap1, Tap2, Psmb8, Psmb9,* and *Nlrc5* were decreased in E cancer cells located closer to qM cancer cells **(Fig 3C, S6C)**. These findings suggest that qM cancer cells can induce MHC-I downregulation in neighboring E cancer cells to generate gradients of immune-evasion and promote cross-protection.

**Figure 3.**
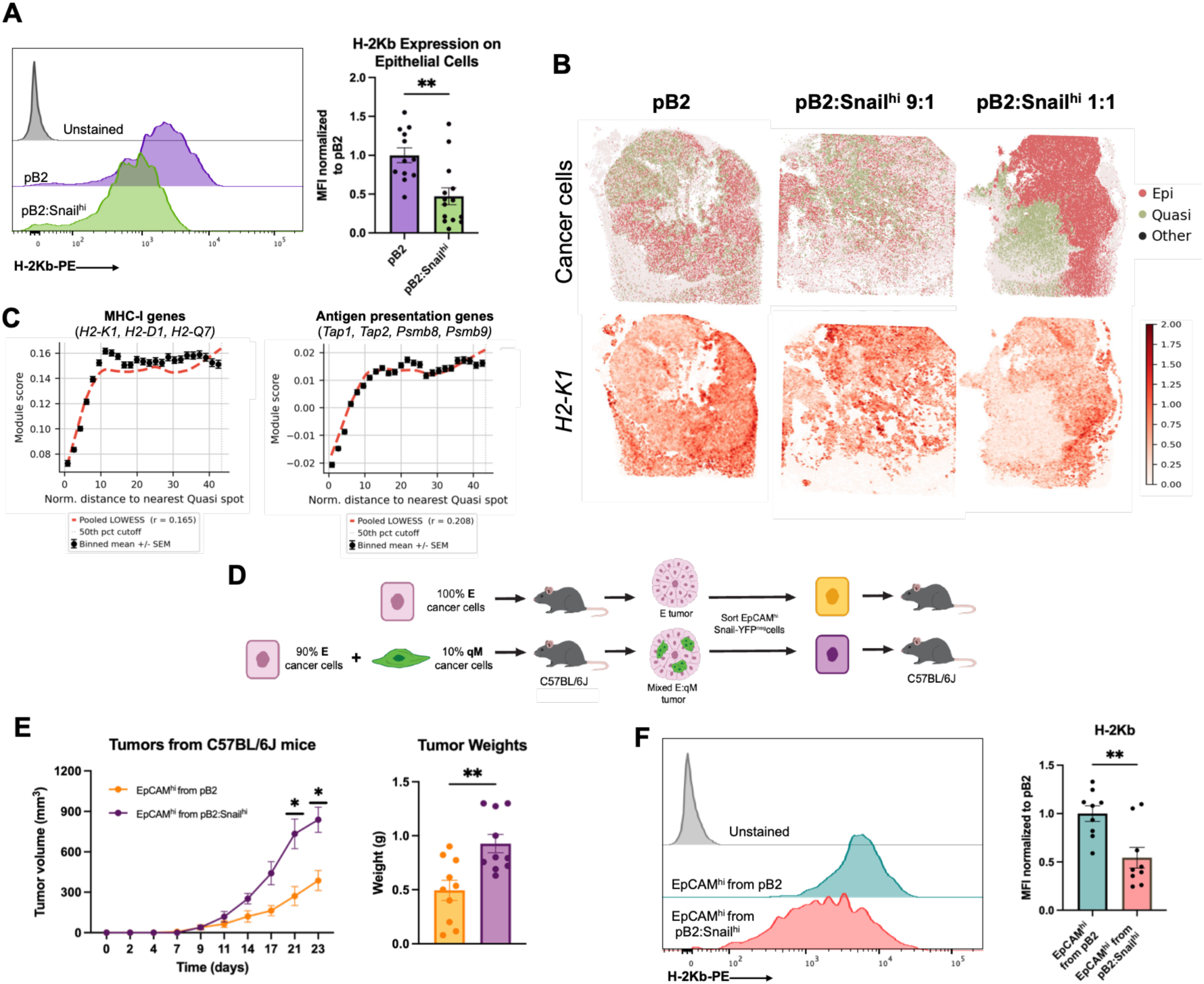
Epithelial cancer cells display MHC-I downregulation within mixed tumors. **(A)** H-2Kb expression measured on CD45^neg^ Snail-YFP^neg^ EpCAM^hi^ epithelial cancer cells in pB2 and pB2:Snail^hi^ tumors, expressed as median fluorescent intensity (MFI) fold-change relative to pB2 tumors with representative histograms. **(B)** Spatial distribution of epithelial and quasi-mesenchymal cancer cells along with *H2-K1* expression in epithelial and mixed 9:1 and 1:1 tumors. **(C)** MHC class I (left; H2-K1, H2-D1, H2-Q7) and antigen presentation (right; Tap1, Tap2, Psmb8, Psmb9, Nlrc5) module scores in epithelial spots versus normalized distance to the nearest qM spot in pB2:Snail^hi^ 1:1 tumor. Points, binned means +/- SEM; dashed line, LOWESS fit to epithelial spots pooled across sections; r, Spearman correlation across pooled epithelial spots. **(D)** Schematic for isolation of epithelial cells from pB2 and pB2:Snail^hi^ tumors and reimplantation into syngeneic hosts. **(E)** Tumor kinetics and weights of epithelial cells isolated from pB2 (orange) and pB2:Snail^hi^ tumors (purple) grown in C57BL/6J mice. **(F)** H-2Kb MFI on epithelial cancer cells from tumors in E, normalized to cancer cells from pB2 tumor. **A** represents four independent experimental replicates including *n*=3-5 per group. **B-C** represent one experimental replicate per condition. **E-F** include three independent experimental replicates with *n*=3-4 per group. Data in **A** and **E-F** are represented as mean ± standard error of the mean (SEM) and were analyzed using a mixed effects model (tumor kinetics, **E**) or unpaired two-tailed t-test (MFI values from flow cytometry, **A** and **F**; tumor weights, **E**). \*\**p* < 0.01. Comparisons are not significant (ns, *p* ≥ 0.05) if not indicated.

We next wanted to understand whether cross-protective effects are transient and require physical juxtaposition of epithelial and quasi-mesenchymal cancer cells. Accordingly, we implanted epithelial pB2 and mixed pB2:Snail^hi^ tumors into the mammary fat pad of C57BL/6J mice. We then used fluorescence-activated cell sorting (FACS) to isolate CD45^neg^ EpCAM^hi^ Snail-YFP^neg^ E cancer cells from both pB2 and pB2:Snail^hi^ tumors. We reimplanted these cells into the mammary fat pad of C57BL/6J mice and calculated tumor kinetics **(Fig 3D)**. We found that epithelial cells isolated from pB2:Snail^hi^ tumors formed larger tumors than those isolated from pB2 tumors grown in C57BL/6J mice **(Fig 3E, S7A).** We confirmed by western blot and RNA sequencing that FACS-isolated epithelial cells do not transition to a mesenchymal state, indicating that the increased tumorigenicity is not due to EMP activation **(Fig S7B-C).** This suggests that epithelial cancer cells retain immune-evasive/tumorigenic properties conferred by quasi-mesenchymal cancer cells, even after physical separation. To determine whether the increased tumorigenicity of epithelial cells derived from mixed tumors could be attributed to MHC-I downregulation, we measured H-2Kb expression on the reimplanted tumors by flow cytometry. Indeed, we found that epithelial cells originating from mixed tumors retained decreased MHC-I levels relative to epithelial tumors, even upon separation from qM cancer cells and reimplantation into new hosts. **(Fig 3F, S7D).**

Since MHC-I is an inhibitory molecule for NK cells^61,62^, we wanted to determine if MHC-I downregulation sensitizes mixed tumors to attack by NK cells. We first visualized the location of NK cells within our Slide-seq samples. We found that NK cells infiltrated all three tumors and co-localized to E regions in mixed tumors **(Fig S7E)**. To determine whether NK cells are functionally important in the control of mixed tumors, we first implanted mixed pB2:Snail^hi^ tumors in NK-proficient C57BL/6J and *Rag1*^-/-^ mice as well as NK cell-deficient NSG mice. Although we did not find a difference in tumor kinetics between these three hosts, pB2:Snail^hi^ tumors implanted in NSG mice had greater tumor weights than those implanted in C57BL/6J and *Rag1*^-/-^ mice **(S7F)**. To investigate this further, we depleted NK cells from mixed pB2:Snail^hi^-tumor-bearing *Rag1*^-/-^ mice using anti-NK1.1 treatment. We found that neither tumor size nor weight was significantly altered upon treatment with anti-NK1.1 **(Fig S7G)**. Together, this suggests that NK cells likely do not play a significant role in controlling mixed tumor growth.

### Exclusion and dysfunction of CD8^+^ T-cells mediate E cancer cell-intrinsic MHC-I downregulation in mixed tumors

Previously, we investigated the location of T-cells within mixed tumors by immunofluorescent staining and found that E regions, but not qM regions, of mixed tumors recruited T-cells^43^. We next asked whether there are differences in the quantity of T-cells recruited to E regions of mixed tumors compared to completely E tumors. To this end, we performed immunofluorescent staining for CD3 and Snail-YFP (to demarcate E and qM regions within mixed tumors) **(Fig 4A-B)**. We found that the total number of CD3^+^ T-cells was decreased in pB2:Snail^hi^ tumors compared to pB2 tumors **(Fig S8A)**. Within pB2:Snail^hi^ tumors, qM regions recruited fewer T-cells compared to E regions **(Fig 4A-C, S8B)**. Finally, there were also fewer T-cells at the periphery of pB2:Snail^hi^ tumors compared to pB2 tumors **(Fig 4A-C)**.

**Figure 4.**
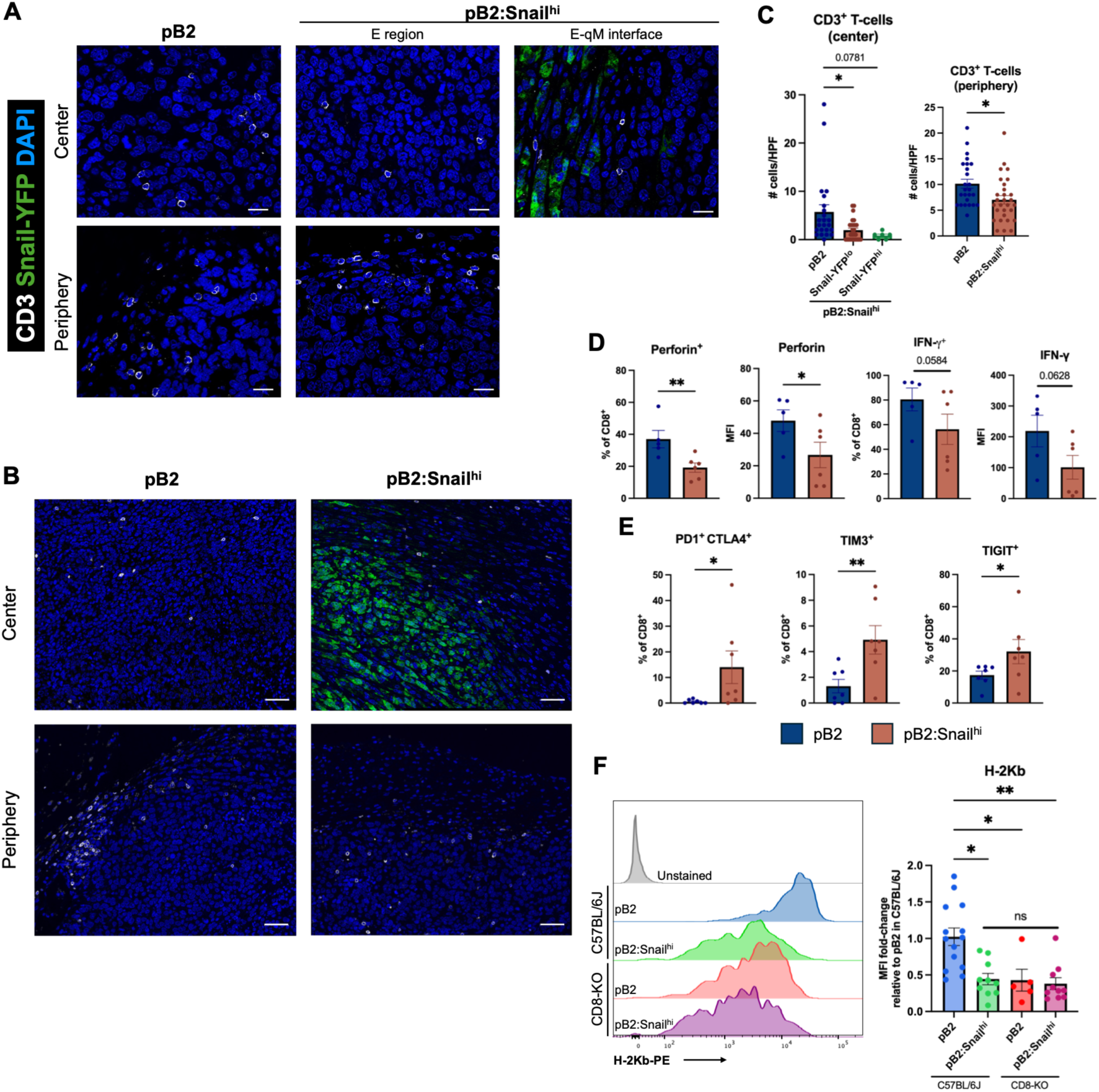
Intratumoral CD8^+^ T-cells regulate epithelial cancer cell-intrinsic MHC-I expression. **(A)** Representative images of immunofluorescent staining for CD3 (white), Snail-YFP (green) and DAPI (blue) in pB2 and pB2:Snail^hi^ tumors. Scale bars indicate 20 μm. **(B)** Tiled images of CD3 staining in pB2 and pB2:Snail^hi^ tumors. Scale bars indicate 50 μm. **(C)** Quantification of CD3^+^ T-cells at the tumor center and periphery across five fields of view in five tumors. **(D)** Flow cytometry data demonstrating percentage of CD8^+^ T-cells expressing activation markers and median fluorescent intensity (MFI) in epithelial and mixed 9:1 tumors. **(E)** Percentage of CD8^+^ T-cells expressing dysfunction markers in epithelial and mixed tumors. **(F)** H-2Kb MFI measured on CD45^neg^ Snail-YFP^neg^ EpCAM^hi^ epithelial cancer cells in pB2 and pB2:Snail^hi^ tumors grown in C57BL/6J or CD8-KO mice with representative histograms. **A-C** represent two independent experimental replicates including *n*=5 total per group. **D-E** include two independent experimental replicates with *n*=3-5 per group. **F** includes two or three independent experimental replicates with *n*=2-5 per group. Data in **C-F** are represented as mean ± standard error of the mean (SEM) and were analyzed using a one-way ANOVA (**C,** left), an unpaired two-tailed t-test **(C,** right), a one-tailed Mann-Whitney test **D-E)**, or a Kruskal-Wallis test (**F**). \**p* < 0.05, \*\**p* < 0.01. Comparisons are not significant (ns, *p* ≥ 0.05) if not indicated.

To determine whether intratumoral CD8^+^ T-cells functionally differ between pB2 and pB2:Snail^hi^ tumors, we measured markers of activation and dysfunction by flow cytometry. We found that CD8^+^ T-cells showed decreased perforin and TNF-α expression, as well as a trend towards decreased IFN-γ expression in mixed tumors **(Fig 4D, S8C)**. CD8^+^ T-cells within mixed tumors showed a concomitant increase in expression of dysfunction markers PD-1, CTLA-4, TIM3, and TIGIT **(Fig 4E)**. These findings are consistent with previously published findings in mixed tumors of different ratios^43^. Together, these results suggest that minority fractions of qM cancer cells within mixed tumors lead to reduced T-cell infiltration and greater T-cell dysfunction compared to CD8^+^ T-cells in E tumors.

To ascertain whether intratumoral CD8^+^ T-cells promote MHC-I expression in E tumors, we measured H-2Kb on E cancer cells within E and mixed tumors implanted in C57BL/6J and CD8-KO mice. We found that compared to E tumors in C57BL/6J mice, E cancer cells within mixed tumors in C57BL/6J mice as well as both E and mixed tumors in CD8-KO showed decreased H-2Kb expression by MFI **(Fig 4F, S8D)**. This suggests that tumor-infiltrating CD8^+^ T-cells in E tumors are necessary to promote cancer cell-intrinsic MHC-I expression. Along these lines, reduced CD8^+^ T-cell infiltration and a dysfunctional phenotype within mixed tumors likely contribute to reduced E cancer cell-intrinsic MHC-I expression.

### Intratumoral macrophage recruitment contributes to mixed tumor growth

To determine if MHC-I downregulation can also occur independently of lymphocytes, we repeated the epithelial cell reimplantation experiment **(Fig 3D)** using pB2 and pB2:Snail^hi^ tumors grown in immunodeficient NSG mice. We found that mixed pB2:Snail^hi^ tumors grew faster than pB2 tumors in NSG mice **(Fig S8E-F).** Epithelial cells isolated from pB2:Snail^hi^ tumors showed a trend towards increased growth compared to those isolated from pB2 tumors grown in NSG mice, but the difference was not significant **(Fig S8G-H)**. We then measured H-2Kb expression on reimplanted E cancer cells from pB2 and pB2:Snail^hi^ tumors grown in NSG mice. Like the tumors grown in CD8-KO mice, E cancer cells reimplanted from tumors in NSG mice showed significantly decreased H-2Kb expression compared to E cancer cells originating from pB2 tumors grown in C57BL/6J mice **(S7D)**. However, in the case of NSG-derived tumors, E cancer cells isolated from pB2:Snail^hi^ tumors showed decreased H-2Kb compared to those from pB2 tumors **(Fig 5A)**. This suggests that in addition to CD8^+^ T-cells, myeloid cells and/or qM-derived paracrine factors may also contribute to decreased MHC-I on E cancer cells in mixed tumors.

**Figure 5.**
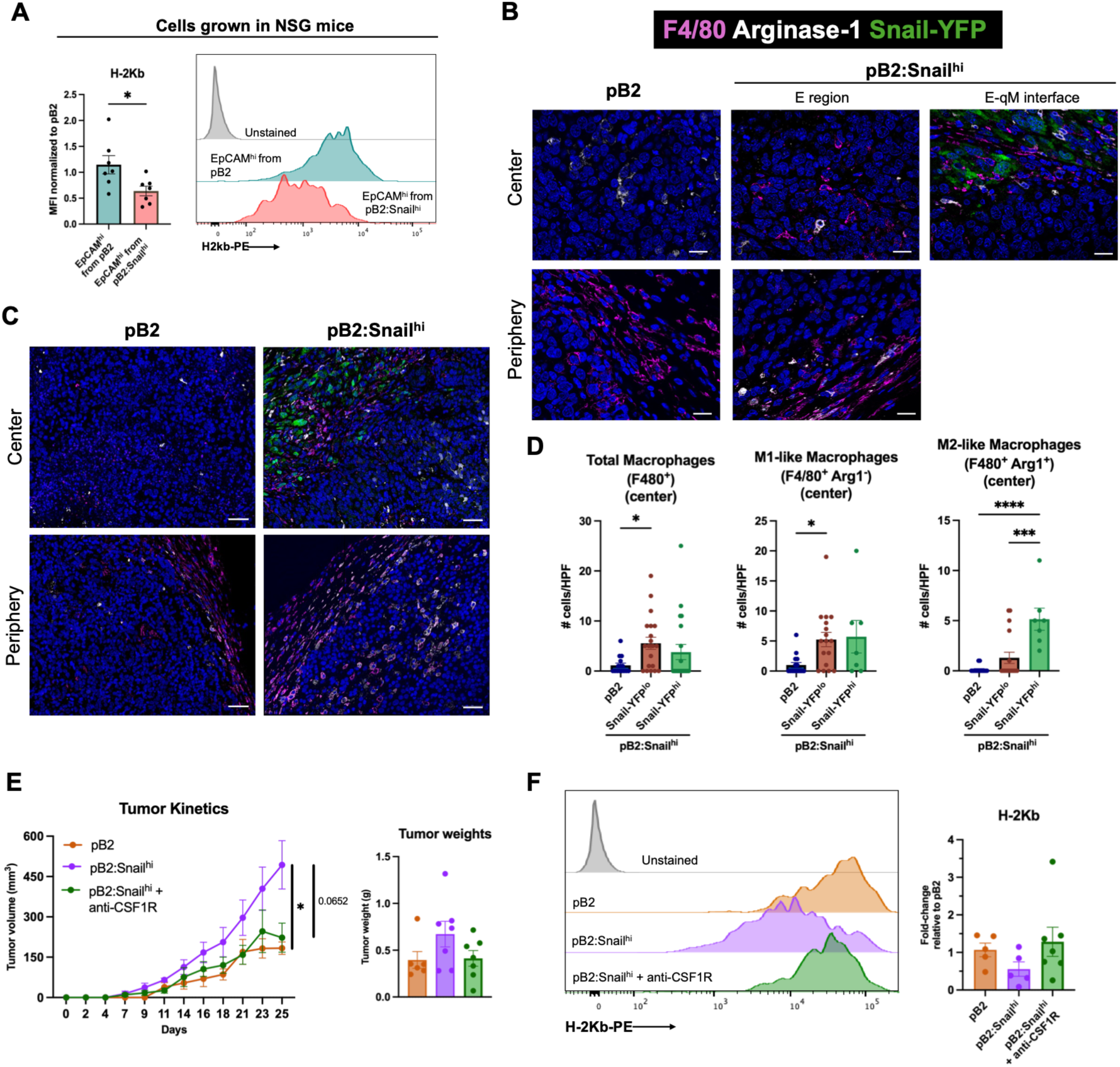
Depletion of intratumoral macrophages reduces mixed tumor growth. **(A)** H-2Kb MFI on epithelial cancer cells isolated from pB2 or pB2:Snail^hi^ tumors grown in NSG and reimplanted in C57BL/6J mice. **(B)** Representative images of immunofluorescent staining for F4/80 (magenta), arginase-1 (white), Snail-YFP (green), and DAPI (blue) in pB2 and pB2:Snail^hi^ tumors. Scale bars indicate 20 μm. **(C)** Tiled images of F4/80 and Arginase-1 (Arg1) staining in pB2 and pB2:Snail^hi^ tumors. Scale bars indicate 50 μm. **(D)** Quantification of tumor-infiltrating Arg1^-^ M1-like macrophages and Arg1^+^ M2-like macrophages across five fields of view in five tumors. **(E)** Tumor kinetics and weights of pB2 and pB2:Snail^hi^ tumors with macrophage depletion. **(F)** H-2Kb MFI on epithelial cancer cells in epithelial or mixed tumors with or without macrophage depletion, normalized to pB2 tumors. **A** includes two independent experimental replicates with *n*=3-4 per group. **B-D** represent two independent experimental replicates including *n*=5 total per group. **E-F** include two independent experimental replicates with *n*=2-4 per group. Data in **A and D-F** are represented as mean ± standard error of the mean (SEM) and were analyzed using an unpaired two-tailed t-test (**A**), one-way ANOVA (**D**), mixed-effects model (tumor kinetics in **E**), or Kruskal-Wallis test (tumor weights in **E** and **F)**. \**p* < 0.05, \*\*\**p* < 0.001, \*\*\*\**p* < 0.0001. Comparisons are not significant (ns, *p* ≥ 0.05) if not indicated.

We first asked whether qM-derived paracrine factors can directly regulate E cancer cell-intrinsic MHC-I expression. We previously demonstrated that qM cancer cells upregulate several immunosuppressive factors^42^. Among these are CD73 and TGFβ-1, both of which are implicated in MHC-I downregulation. Specifically, CD73-derived adenosine has been shown to reduce MHC-I levels on dendritic cells and TGFβ-1 downregulates MHC-I in multiple tissue types^63–67^. To determine whether these qM-derived factors mediate E cancer cell-intrinsic MHC-I downregulation, we generated mixed tumors where the qM cancer cells were knocked out for either CD73 or TGFβ-1. E cancer cells within pB2:sgCD73 and pB2:sgTGFB1 mixed tumors retained decreased MHC-I expression compared to E tumors, suggesting that qM-derived adenosine and TGFβ-1 do not significantly contribute to MHC-I downregulation **(Fig S9A)**.

After determining that qM-derived adenosine and TGFβ-1 do not directly regulate E cancer cell-intrinsic MHC-I, we next investigated the involvement of TAMs. Previously, we found that qM regions within mixed tumors recruit increased Arginase-1 (Arg1)^+^ M2-like macrophages compared to E regions^43^. We first tested whether the quantity of tumor-infiltrating macrophages differs between pB2 and pB2:Snail^hi^ tumors. By immunofluorescent staining, we found that pB2:Snail^hi^ tumors recruit increased M1-like (F4/80^+^ Arg1^-^) and M2-like (F4/80^+^ Arg1^+^) macrophages compared to pB2 tumors **(Fig 5B-D, S9B-C)**. There was an equal distribution of M1-like macrophages between E and qM regions within pB2:Snail^hi^ tumors, whereas M2-like macrophages were preferentially recruited to qM regions **(Fig 5D, S9B-C).** We did not find significant differences in the number of M1-like or M2-like macrophages at the periphery of pB2 and pB2:Snail^hi^ tumors **(Fig S9C).** The percentage of CD206^+^ M2-like macrophages was also increased in pB2:Snail^hi^ tumors by flow cytometry, in accordance with previous findings^43^ **(Fig S9D)**.

To determine whether the increased influx of macrophages promotes mixed tumor progression, we depleted macrophages from mixed tumor-bearing mice using anti-CSF1R treatment **(Fig S9E-F)**. We found that macrophage-depleted mice formed smaller mixed tumors, comparable in size to completely E tumors **(Fig 5E, S9G)**. By flow cytometry, E cancer cells in macrophage-depleted mixed tumors showed a trend towards restored H-2Kb expression, similar in levels to pB2 tumors **(Fig 5F, S9H)**. This suggests that both CD8^+^ T-cells and macrophages may control E cancer cell-intrinsic MHC-I expression in mixed tumors.

### Human triple-negative breast cancers display EMP-associated T-cell exclusion

To assess the translational relevance of our findings, we used publicly-available human TNBC spatial transcriptomics data^68^ to investigate associations between EMP activation and immune cell infiltration. First, we visualized enrichment of the Molecular Signature Database HALLMARK_EPITHELIAL_MESENCHYMAL_TRANSITION^69^ gene set in tumor cells **(Fig 6A)**. We found that distinct regions show enrichment of this signature, which is modeled by our murine mixed tumor model. Next, we asked whether TNBC samples with differential EMP activation showed differences in T-cell infiltration. We found that samples with low EMP activation displayed infiltration of effector CD8^+^ T-cells, whereas effector CD8^+^ T-cells were restricted to the surrounding stroma in samples with high EMP activation **(Fig 6B, S10A).** We observed a similar pattern in exhausted CD8^+^ T-cells, but with reduced numbers of cells in some samples with low EMP activation **(Fig 6C, S10B)**.

**Figure 6.**
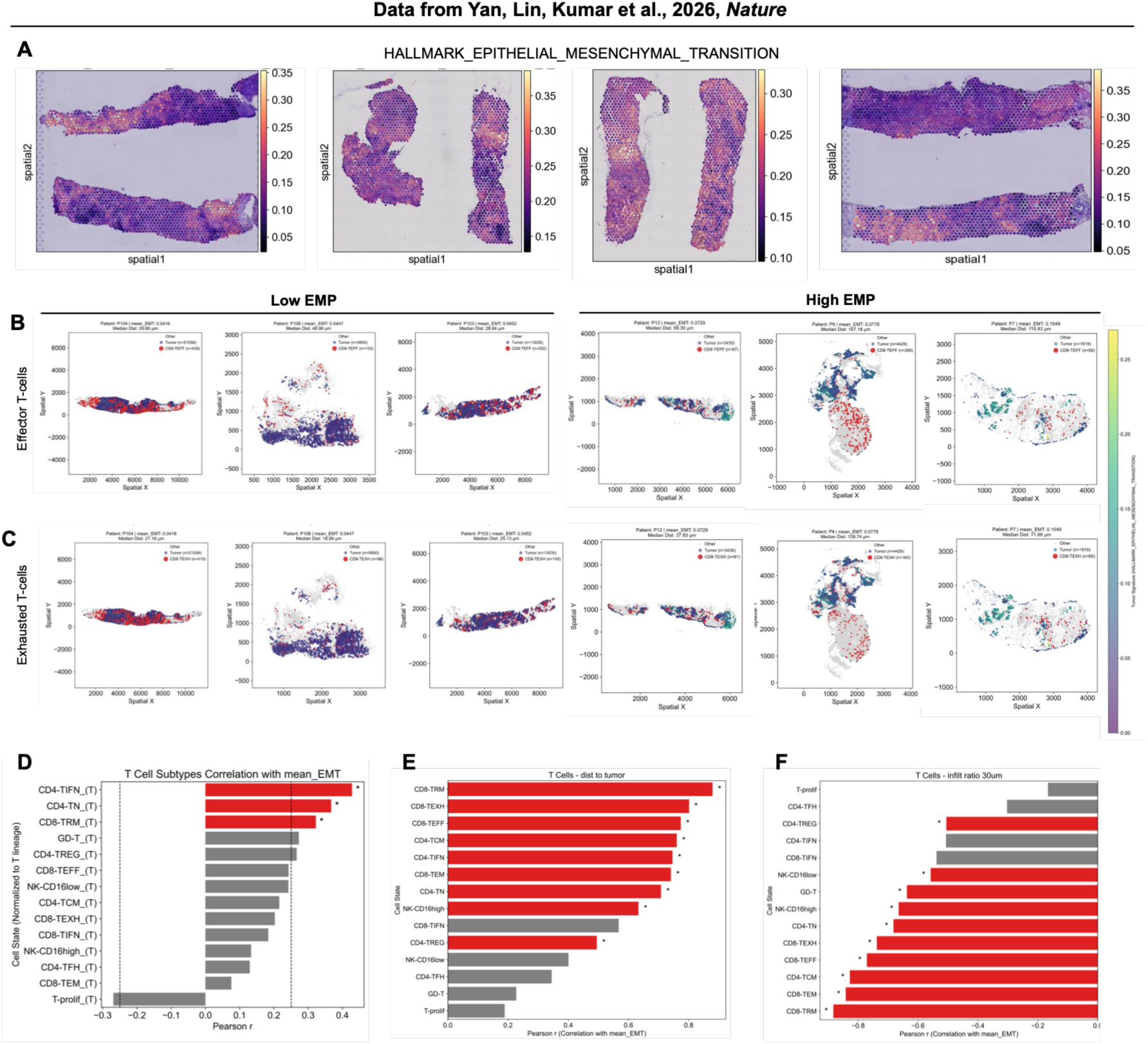
Epithelial-mesenchymal plasticity is associated with T-cell exclusion in human triple-negative breast cancers. **(A)** Expression of the Molecular Signatures Database HALLMARK_EPITHELIAL_MESENCHYMAL_TRANSITION gene set in triple-negative breast cancer (TNBC) patient samples. **(B)** Distribution of effector CD8^+^ T-cells in TNBC samples with low (left) and high (right) EMP activation. Tumor cells are colored by EMP enrichment and effector CD8^+^ T-cells are depicted in red. **(C)** Distribution of exhausted CD8^+^ T-cells in TNBC samples with low (left) and high (right) EMP activation. Tumor cells are colored by EMP enrichment and exhausted CD8^+^ T-cells are depicted in red. **(D)** Correlation of the proportion of each T-cell subtype with mean epithelial-mesenchymal transition (EMT) score in tumor cells. **(E)** Correlation of the distance of each T-cell subtype from nearest tumor cell with mean epithelial-mesenchymal transition (EMT) score in tumor cells. **(F)** Correlation of the infiltration ratio of each T-cell subtype with mean epithelial-mesenchymal transition (EMT) score in tumor cells. Data from Yan, Lin, Kumar et al., 2026, *Nature.* **D-F** include 44 TNBC patient samples. In **D-F**, red bars and * indicate false discovery rate (FDR) < 0.05.

To ascertain whether these visual differences in T-cell infiltration were significant, we calculated Pearson r correlation coefficients for the presence, distance, and infiltration of T-cells with EMP activation in cancer cells. Using published immune cell states^68^, we found that the proportions of interferon-responsive CD4^+^ T-cells (CD4-TIFN), naïve CD4^+^ T-cells (CD4-TN), and resident memory CD8^+^ T-cells (CD8-TRM) were significantly associated with EMP activation. γδ T-cells (GD-T) and CD4^+^ regulatory T-cells (CD4-TREG) displayed a trend towards increased enrichment in samples with increased EMP activation. Conversely, proliferating T-cells (T-prolif) showed a trend towards reduced enrichment in samples with increased EMP activation **(Fig 6D)**. Next, we assessed the distance of each T-cell subset from the nearest cancer cell. We found that CD8-TRM, exhausted CD8^+^ T-cells (CD8-TEXH), effector CD8^+^ T-cells (CD8-TEFF), central memory CD4^+^ T-cells (CD4-TCM), CD4-TIFN, effector memory CD8^+^ T-cells (CD8-TEM), CD4-TN, CD16^high^ NK cells (NK-CD16high), and CD4-TREG showed significantly increased distance from cancer cells in tumors with increased EMP activation **(Fig 6E)**. Finally, we investigated the infiltration of each T-cell subset within 30 μm of a cancer cell. We found that CD4-TREG, CD16^low^ NK cells (NK-CD16low), GD-T, NK-CD16high, CD4-TN, exhausted CD8-TEXH, CD8-TEFF, CD4-TCM, CD8-TEM, and CD8-TRM showed significantly decreased infiltration in tumors with increased EMP activation **(Fig 6F)**.

Finally, we investigated whether myeloid cells also showed associations in presence, distance, and infiltration in tumors with increased EMP activation. Plasmacytoid dendritic cells (pDC), extracellular matrix remodeling-related macrophages (mac-ECM), and mature dendritic cells (mDC) were significantly enriched in tumors with increased EMP activation. Mac-M2-CXCL macrophages and neutrophils also showed a trend towards enrichment, whereas mac-M1-CCL macrophages and lipid-related mac-lip-C1Q macrophages showed a trend towards decreased presence **(Fig S10C)**. Among all myeloid cell states, only neutrophils displayed a significant association between distance and infiltration with tumors with increased EMP activation **(Fig S10D-E)**. Together, these findings indicate that EMP is associated with T-cell exclusion in human TNBC.

## Discussion

Both EMP and tumor heterogeneity are drivers of treatment resistance in breast cancers. The phenomenon of subclonal cancer cell coexistence and cooperation within breast tumors has been described in the context of genetic heterogeneity and Wnt signaling, but the role of EMP and immunosuppression is poorly understood^70–73^. In this work, we describe a mechanism by which minority fractions of qM cancer cells cross-protect heterogeneous tumors against immune attack through indirect modulation of MHC-I levels on E cancer cells. We first show that cross-protection requires physical proximity of E and qM cancer cells, rather than acting systemically. Next, we identify that minority fractions of qM cancer cells prevent CD8^+^ T-cells from constraining mixed tumor growth through altered location and function. Additionally, E cancer cells become resistant to attack by CD8^+^ T-cells through long-term MHC-I downregulation. We next investigated the mechanism of this immune-evasion and found that both CD8^+^ T-cells and macrophages contribute to MHC-I regulation in mixed tumors. Finally, we found that EMP activation is associated with exclusion of multiple CD8^+^ and CD4^+^ T-cell subsets in human TNBC patient samples. These results identify qM cancer cells as indirect downregulators of E cancer cell-intrinsic MHC-I through altered T-cell and macrophage infiltration and function in mixed tumors.

Tumor development can induce systemic immunosuppression, for example through hemopoietic dysregulation resulting in reduced frequencies of CD4^+^ and CD8^+^ T-cells and increased production of immunosuppressive myeloid populations and Tregs^74,75^. T-cells in tumor-bearing mice also incur functional deficits including reduced IFN-γ production^74,76^. Therefore, we first aimed to determine whether cross-protection occurs due to qM-mediated systemic immunosuppression. We found that E tumors did not become immunosuppressive upon contralateral implantation of a qM tumor. This is consistent with studies on myeloid cell infiltration in a distinct contralateral model^77^. Moreover, circulating immune cells were not significantly altered by the presence of a qM tumor in our model. Because ICB therapeutic efficacy also relies on intact peripheral immunity, we also tested the sensitivity of E tumors to anti-CTLA-4 treatment in the presence of a contralateral qM tumor. E tumors also remained sensitive to anti-CTLA-4 therapy in the presence of a qM tumor, suggesting that systemic immunity is not impaired by qM tumors. However, we did not test whether qM-derived immunosuppressive factors were actually in circulation. Nonetheless, based on the lack of immunosuppression induced by contralateral qM tumors, we decided to focus on immunosuppressive changes within the local TME of heterogeneous tumors.

Next, we sought to understand changes to the location and function of immune cells within mixed tumors. We found that unlike E tumors, mixed tumors grew at similar rates in immunocompetent and CD8-KO mice, suggesting that qM cancer cells prevent CD8^+^ T-cells from constraining mixed tumors. By spatial transcriptomics, we identified that qM cancer cells create distinct immunosuppressive niches within heterogeneous tumors. This finding suggests that E and qM cancer cells may express distinct chemokines to control recruitment of immune cells. Moreover, immunosuppressive TAMs recruited to qM regions could inhibit the antitumor activity of T-cells within E regions. These results are in alignment with emerging insights from spatially resolved technologies that demonstrate co-localization of immunosuppressive cells and cytokines with malignant, EMP-activated cells^78–81^. In fact, EMP-enriched cancer cells have been shown to co-localize with SPP1^+^ TAMs in human cisplatin-resistant muscle-invasive bladder cancer^82^. Similarly, metastasis-initiating hybrid E/M cancer cells have been shown to spatially overlap with TAMs in TNBC^83^. Finally, CD8^+^ T-cells within mixed tumors upregulated dysfunction markers. This may be attributed to immunosuppressive factors expressed by qM cancer cells, such as CD73 and TGF-β1^42^.

We found that E cancer cells also become resistant to control by CD8^+^ T-cells through long-term downregulation of MHC-I. MHC-I is expressed ubiquitously on the cell surface and is essential for antigen presentation to CD8^+^ T-cells. Expression of MHC-I is tightly regulated by epigenetic modifications and transcription factors including NF-κB and NLRC5. Most cells constitutively express low levels of MHC-I, but upregulate genes required for antigen presentation on MHC-I in response to interferons (IFN)^84,85^. Because cancer-associated or -specific antigens presented on MHC-I allow CD8^+^ T-cells to identify and kill cancer cells, a common mechanism of immune evasion is MHC-I loss^84–87^. Cancer cell-intrinsic MHC-I downregulation has implications for cancer immunosurveillance, as well as immunotherapy efficacy. Several chemokines are also induced by IFN-γ^88,89^, which could suggest that reduced CD8+ T-cell-derived IFN-γ within mixed tumors leads to downregulation of MHC-I and T-cell-attracting chemokines.

EMP has previously been linked to MHC-I reduction in quasi-mesenchymal cancer cells. EMP-TFs including Snail and Zeb1 directly repress genes involved in antigen processing and presentation^43,90–92^. Moreover, the receptor tyrosine kinase AXL is linked with EMP and regulates MHC-I expression^93^. Here, we demonstrate a distinct mechanism of EMP-mediated MHC-I downregulation where cancer cells remain epithelial while downregulating MHC-I in response to neighboring qM cancer cells. Our results reveal that MHC-I regulation on E cancer cells can be driven by qM cancer cells within heterogeneous tumors through T-cell exclusion and macrophage recruitment. These findings can provide therapeutic targets for reversing MHC-I downregulation in heterogeneous tumors, for example by targeting immunosuppressive qM-derived factors or using agents that promote T-cell infiltration or restrict macrophage infiltration.

Cancer cell-intrinsic MHC-I expression has been previously linked to TAMs. Specifically, engagement of the MHC-I component β2-microglobulin on cancer cells by LILRB1 on macrophages prevents cancer cell phagocytosis^94^. Previous work has also demonstrated a role for TAMs in controlling MHC-I-deficient tumors through acquisition of a M1-like phenotype and increased antigen cross-presentation^95^. These findings suggest that TAMs not only regulate E cancer cell-intrinsic MHC-I expression, but MHC-I downregulation may also sensitize mixed tumors to phagocytosis by macrophages.

Several questions remain from the results presented here. First, the downregulation of MHC-I would be expected to sensitize E cancer cells in mixed tumors to attack by NK cells. Implanting mixed tumors in NSG mice resulted in greater tumor weights compared in mixed tumors grown in NK-proficient C57BL/6J or *Rag1*^-/-^ mice. However, we found that depletion of NK cells did not alter mixed tumor growth. This may be due to the relatively small proportion of NK cells within mixed tumors **(Fig S5C)**. It is also unknown whether E cancer cells express NK cell-activating ligands. Finally, the NK cells present in mixed tumors may be dysfunctional due to the immunosuppressive factors secreted by qM cancer cells (e.g. TGF-β1, adenosine).

The finding that E cancer cells isolated from mixed tumors maintain reduced MHC-I upon reimplantation into new hosts suggests an epigenetic component to this mechanism of cross-protection. Alternatively, qM residence in mixed tumors may downregulate chemokine expression by E cancer cells, preventing T-cell infiltration in reimplanted tumors. Previous studies have demonstrated that MHC-I-deficient tumors exhibit immune desertification, suggesting this could be a mechanism of long-term MHC-I downregulation^95^. Finally, it is unknown how qM cancer cells control T-cell and macrophage infiltration in mixed tumors. Moreover, although we have identified macrophages as a regulator of MHC-I expression, specific macrophage-derived factors that control MHC-I expression remain to be uncovered.

Nonetheless, these findings highlight minority fractions of qM cancer cells as indirect regulators of E cancer cell-intrinsic MHC-I expression. The mixed tumor model used here is translationally relevant, as it recapitulates the heterogeneity in epithelial-mesenchymal cell states present in spontaneous breast tumors. By performing spatial transcriptomics on our mixed tumors, we identified that tumor-infiltrating immune cells create niches within epithelial and quasi-mesenchymal regions of mixed tumors. These niches, as well as functional impairments in CD8^+^ T-cells lead to decreased MHC-I on E cancer cells. We have also identified a role for tumor-associated macrophages in downregulating cancer cell-intrinsic MHC-I. These results demonstrate a mechanism of immune evasion in heterogeneous breast tumors, whereby minority fractions of qM cancer cells alter immune infiltration and function, leading to E cancer cell-intrinsic MHC-I downregulation.

## Materials and Methods

### Mice

C57BL/6J, *Rag1*^-/-^, and *Cd8a*^tm1Mak^ (CD8-KO) female mice, aged 6–8-weeks, were obtained from The Jackson Laboratory. NOD.Cg-*Prkdc^scid^ Il2rg^tm1Wjl^*/SzJ (NSG) mice were accessed through the Cornell University Progressive Advancement of Therapeutics (PATh) facility. Mice were age matched and randomly assigned to treatment or control groups in all experiments. All animal procedures were carried out in compliance with guidelines and protocols approved by the Institutional Animal Care and Use Committee (IACUC) under protocols 2021-0115 and 2017-0035 and maintained by the Center for Animal Resources and Education (CARE) at Cornell University.

### Cell lines and tissue culture

Murine cell lines pB2, Snail^hi^, EpCAM^hi^, sgCD73, and sgTGFβ1 were a kind gift from the Weinberg Lab. These cell lines were maintained as previously described^42,43,57^. Murine cell lines were cultured in a 1:1 mixture of Dulbecco’s modified Eagle’s medium (DMEM) and Ham’s F12 medium supplemented with 5% bovine adult serum, 1x penicillin-streptomycin and 1x nonessential amino acids (hereafter referred to as MMTV-PyMT media) maintained in a 37°C incubator containing 5% CO2. All cell lines were routinely tested for mycoplasma (from 2019-2025) using the MycoAlert Mycoplasma Detection Kit (Lonza). All cell lines are negative for Mycoplasma.

### In vivo mouse models and tissue dissociation

For orthotopic tumor implantations, 1 × 10^6^ cells were counted and resuspended in 30 μL of media containing 20% Matrigel. Cells were then implanted orthotopically into the mammary fat pads of C57BL/6J mice. Tumor volume was calculated using the modified ellipsoid formula tumor volume (mm^3^) = *(L x W x W)/*2, where *L* represents the largest tumor diameter and *W* represents the perpendicular measurement. After the tumors reached 2000 mm^3^, mice were euthanized and tumors were collected. A small section of the tumors was saved for fixing while the rest was used for making single-cell suspensions for flow cytometry. For tumor digestions, tumors were cut and finely minced with a razor blade and digested in RPMI containing 2mg/ml Collagenase A (Krackeler Scientific) and 100 U/ml Hyaluronidase (Krackeler Scientific). The suspension was then incubated in a rotator at 37°C for 40 mins. After digestion, the single-cell suspension was filtered first through a 70-micron and then through a 40-micron pore sized strainer, followed by centrifugation at 1250rpm for 10 mins at 4°C. The pellets were then resuspended in RPMI containing Monensin GolgiStop (BD Sciences) and incubated for 3-5 hours at 37°C. After incubation, the cells were centrifuged at 1250rpm for 10 mins at 4°C and processed for flow cytometry as described below.

### Flow cytometry of tumors

The following steps were performed in 96-well V-bottom microwell plates using single cell suspensions obtained after tumor processing. First, cells were centrifuged at 1250rpm for 5 mins at 4°C and the pellets were resuspended in 200ul of FACS wash buffer with a master mix containing surface markers each diluted 1:100: EpCAM PECy7 (G8.8, Invitrogen, RRID: AB_1724047), CD45 BUV563 (30-F11, BD Biosciences, RRID: AB_2870209), CD45 BV421 (30-F11, Invitrogen, RRID: AB_2562559), H2kb PE (AF6-88.5.5.3, eBioscience, RRID: AB_10598797), CD8a APC (53-6.7, Invitrogen, RRID:AB_469335), PD1 FITC (J43, Invitrogen, RRID: AB_465472), CTLA4 PE (UC10-4B9, Invitrogen, RRID: AB_465879), TIM3 PerCP-Cyanine5.5 (RMT 3-23, BioLegend, RRID: AB_2571934), TIGIT PECy7 (1G9, BioLegend, RRID: AB_2565648), CD11b PerCP-Cyanine5.5 (M1/70, Invitrogen, RRID: AB_953558), F4/80 PECy7 (BM8, BioLegend, RRID: AB_893490), Ly6C BV650 (HK 1.4, BioLegend, RRID: AB_2800630), Ly6G FITC (1ALy6g, Invitrogen, RRID: AB_2572532), I-A/I-E (MHC-II) PE (M5/114.15.2, BioLegend, RRID: AB_313322), CD206 APC (C068C2, BioLegend, RRID: AB_10896057).

Live/Dead staining was performed for 15 mins in the dark at 4°C using the Invitrogen LIVE/DEAD Fixable Near-IR Dead Cell Stain Kit with APC-CY7 dye. Cells were then fixed for 30-60 mins using the Intracellular Fixation and Permeabilization Buffer Set (ThermoFisher). Intracellular staining was performed for 30-60 mins in dark on ice or at 4°C using the FOXP3/Transcription Factor Staining Buffer Set (Thermo Fisher) for the following antibodies each diluted at 1:50: IFN-γ PECy7 (XMG1.2, BioLegend, RRID: AB_1595591), TNF-α BV650 (MP6-XT22, BioLegend, RRID: AB_2562450), Perforin PE (eBioOMAK-D, Invitrogen, RRID: AB_466243). The single stain controls were made using UltraComp eBeads Compensation Beads (Invitrogen). Flow cytometry data were acquired on a BD Biosciences FACSymphony A3, and data were analyzed using the FlowJo (v10, TreeStar) software.

### Epithelial cell reimplantation experiments

To obtain epithelial cells from orthotopically implanted epithelial and mixed tumors in C57BL/6J and NSG mice, tumors were digested as described above. Cells were resuspended in 200 μL of FACS wash buffer with a master mix containing surface markers each diluted 1:100: EpCAM PECy7 (G8.8, Invitrogen, RRID: AB_1724047) and CD45 BV421 (30-F11, Invitrogen, RRID: AB_2562559). The Snail^hi^ quasi-mesenchymal cells contain an endogenous fluorescent Snail-YFP reporter. CD45^neg^ Snail-YFP^neg^ EpCAM^hi^ epithelial cells were sorted from pB2 and mixed pB2:Snail^hi^ tumors into 1 mL MMTV-PyMT media using a BD FACSMelody Cell Sorter. Cells were then plated in 10 cm plates and expanded until enough cells were available for reimplantation (∼1 week). Cells were then orthotopically implanted into the mammary fat pads of C57BL/6J mice as described above. EMP state of the cells was confirmed prior to reimplantation by western blot **(Fig S7B)**.

### In vivo immune checkpoint blockade therapy and immune cell depletion

Mice received 200 μg of anti-CTLA4 (clone 9H10, BioXCell BE0131, RRID: AB_10950184) diluted in 200 μL of PBS and administered intraperitoneally 3x per week for 6 doses followed by 100 μg weekly injections for 2 additional doses. To achieve depletion of NK cells, 300 μg anti-NK1.1 (clone PK136, BioXCell BE0036, RRID: AB_1107737) was administered weekly, starting 3 days prior to tumor cell implantation. To deplete macrophages, 500 μg anti-CSF1R (clone AFS98, BioXCell BE0213, RRID: AB_2687699) was administered 3x per week, starting 7 days prior to tumor cell implantation.

### Immunofluorescence staining of tumor sections

Immunofluorescence staining was performed as previously described^42,43,57^. Tumor sections were fixed in 10% neutral buffered formalin for 24-48 hours and transferred to freshly made 70% ethanol, followed by embedding and sectioning. Formalin-fixed paraffin-embedded (FFPE) tumor sections were deparaffinized in Histoclear and rehydrated through a series of decreasing ethanol concentrations (100%, 95%, 75%, Milli-Q water and 1x DAKO wash buffer; 5 mins each). For antigen retrieval, the section slides were immersed in 1X DAKO antigen retrieval buffer pH 6.0 (Agilent Technologies) and microwaved. Following this, the slides were washed twice with 1x DAKO wash Buffer (Agilent Technologies). To block non-specific binding, sections were blocked with phosphate-buffered saline containing 0.3% TritonX 100 (Millipore Sigma) and 1% Normal donkey serum (Jackson Immunoresearch Lab) for 20 mins at Room temperature (RT). After blocking, the sections were stained overnight at 4°C with anti-E-cadherin (BD Pharmingen 610182, RRID: AB_397581, 1:50), anti-Vimentin (CST 5741S, RRID: AB_10695459, 1:50), anti-CD3 (Abcam ab16669, RRID: AB_443425, 1:100), anti-F4/80 (ThermoFisher MF48000, RRID: AB_10376289, 1:50), anti-Arginase 1 (Santa Cruz Biotechnology sc271430, RRID: AB_10648473, 1:50). The sections were then washed twice with 1X DAKO wash buffer and tagged with Biotium CF555 Donkey anti-Rabbit IgG (RRID: AB_10558011, 1:500) and Biotium CF488A Donkey anti-Mouse IgG (RRID: AB_10561327, 1:500) for 2 hrs. at room temperature. After three washes, the sections were stained with DAPI (1:1000 from 10mg/ml stock-Millipore Sigma) for 5 mins at room temperature. Slides were washed for 5 mins in distilled water and mounted with Prolong Gold anti-Fade mounting media (Cell Signaling Technologies 9071S).

### Microscopy

The stained sections were viewed under a Zeiss AxioObserver inverted microscope with 63X oil objective and the images were analyzed using the ZEN imaging software.

### Western blotting

Lysates were made in aqueous 1X RIPA lysis buffer (EMD Millipore Corp) and the concentration of extracted protein was measured using the Pierce BCA protein assay (Thermo Scientific, 23227) and the optical density was measured on a BioTek Synergy 2 Microplate Reader at 562nm and the data were analyzed in Gen5 1.11 software. 40ug of protein was resolved using the NuPAGE Bis-Tris mini protein gels, 12%,1.0mm (Invitrogen) and transferred to 0.2-micron immobilon-P PVDF membrane (Millipore Sigma) using a wet transfer. The blots were then blocked in 5% Blotto (Santa Cruz) containing 0.2% Tween-20 (Sigma Aldrich) in PBS. Membranes were then probed with primary antibodies (1:1000) overnight, washed and incubated with horseradish peroxidase-labeled secondary antibodies (1:5000) and developed using Dura substrate (Thermo Scientific). Primary antibodies: GAPDH (14C10, Cell Signaling Technologies, RRID: AB_561053), E-Cadherin (24E10, Cell Signaling Technologies, RRID: AB_2291471), N-cadherin (D4R1H, Cell Signaling Technologies, RRID: AB_2687616), Snail (C15D3, Cell Signaling Technologies, RRID: AB_2255011), Zeb1 (D80D3, Cell Signaling Technologies, AB_1904164), Vimentin (D21H3, Cell Signaling Technologies, RRID: AB_10695459) and secondary antibodies: anti-Rabbit IgG, HRP (Cell Signaling Technologies, RRID: AB_2099233) and anti-Mouse IgG, HRP (Cell Signaling Technologies, RRID: AB_330924).

### Tumor processing for single-cell RNA sequencing (scRNA-seq)

Tumor sample processing was performed as previously described^96^. To generate single-cell suspensions, the tumors were first minced using a razor blade and digested in RPMI containing 2 mg/ml Collagenase A (Krackeler Scientific) and 100 U/ml Hyaluronidase (Krackeler Scientific). The suspension was incubated in a rotator at 37°C for 40 mins. Following digestion, the single-cell suspension was filtered through a 70-micron filter, followed by a 40-micron pore-sized strainer. The samples were then centrifuged at 1250 rpm for 10 mins at 4°C. The samples were resuspended in 1 mL of ACK Lysing Buffer (Gibco) and incubated at room temperature for 5 minutes to allow for hypotonic lysis of red blood cells. After 5 minutes, 1 mL warm RPMI was added, and the samples were centrifuged at 1250 rpm for 10 minutes. The pellet was then resuspended in Dead Cell Removal MicroBeads (Miltenyi Biotec) and incubated at room temperature for 15 minutes. After incubation, 500 μL of 1X binding buffer (Miltenyi Biotec) was added and the suspension was placed on a magnet stand for 5 minutes to remove dead cells. The supernatant was then transferred to a new tube and centrifuged at 1250 rpm for 5 minutes. The pellet was resuspended in 6200 μL cold PBS (Gibco) and 1800 μL Debris Removal Solution (Miltenyi Biotec) was mixed with the PBS. Then, 4 mL of cold PBS was overlaid on this mixture, and the tubes were centrifuged at 3000xg for 10 minutes at 4°C. The top two phases were discarded from the tube and 15 mL cold PBS was added to the bottom phase. The tubes were inverted three times and then centrifuged for 10 minutes at 1000xg at 4°C. Finally, the pellet was resuspended in PBS containing 0.04% Bovine Serum Albumin (BSA, Sigma-Aldrich) at a final concentration of 1,000 cells/μl. Single cell suspensions were run on a Chromium X instrument and libraries were prepared following the Chromium GEM-X Single Cell 3’ v4 Assay (10x Genomics, user guide CG000731, RevA) by the Cornell BRC Genomics Facility (RRID:SCR_021727). We targeted 10,000 cells per sample and used 12 cycles of cDNA amplification. Sample quality was confirmed using a Qubit (DNA HS kit; Thermo Fisher) to determine concentration and a Fragment Analyzer (Agilent) to confirm fragment size integrity. Libraries were sequenced on a NovaSeqX, 10B flowcell with 2 x 150bp read length.

### Public scRNA-seq data retrieval and integration with in-house data

Public single-cell and single-nucleus RNA-seq data of mouse mammary tumors were retrieved from the Arc Institute scBaseCount resource (release 2026-01-12, GeneFull, *Mus musculus*). Samples were retained if they were generated with 10x Genomics 3’ gene expression chemistry, annotated as either single-cell or single-nucleus preparations, and annotated as a single disease with high confidence. Of those, samples derived from MMTV-PyMT cell lines and samples with no annotated cell line whose disease annotation corresponded to triple-negative breast cancer were selected for analysis.

Downloaded objects were filtered for cells containing at least 1,000 total counts and merged by accession number as batch covariate. Ensembl gene identifiers were mapped to MGI gene symbols with mygene and made unique. In-house libraries from the three experimental conditions (pB2, pB2:Snail-hi 9:1 and pB2:Snail-hi 1:1) were read from Cell Ranger raw feature-barcode matrices using gene symbols, filtered to cells with at least 1,000 detected genes, and concatenated. Public and in-house matrices were then merged on shared genes, retaining the data source as an additional covariate.

### Quality control, normalization, and integration

Quality metrics were computed with Scanpy (v 1.11.5)^97^, accounting for mitochondrial and ribosomal genes. Cells were retained with fewer than 10% mitochondrial counts, fewer than 40% ribosomal counts, more than 500 detected genes and more than 1,000 total counts. Using Scanpy’s default settings, the counts were normalized and log transformed. The 2,000 most highly variable genes were selected with batch-aware ranking across samples, scaled (values clipped at 10), and reduced by PCA to 75 components. Batch effects across samples were corrected with Harmony (20 iterations), and the corrected embedding was used for all downstream neighbor computations.

### Clustering and cell type annotation

A nearest-neighbor graph was built on 25 Harmony components with 30 neighbors per cell, followed by UMAP embedding. Leiden clustering (igraph implementation) was run with resolution of 0.2, and clusters of interest were subclustered at resolutions of 0.1 to 0.3 to resolve populations that remained mixed at the global resolution. Cluster marker genes were ranked by Wilcoxon rank-sum test, and clusters were assigned to cell identities using canonical markers: Neutrophils (*S100a8, S100a9, Csf3r, Cxcr2*), TAMs (*Mrc1, Arg1, Trem2, Apoe, Folr2*), cDCs (*Xcr1, Clec9a, Flt3, Itgax, Batf3*), Epithelial tumor cells (*Krt8, Krt18, Epcam, Cdh1*), Quasi-mesenchymal tumor cells (*Vim, Nt5e, Twist1, Snai1, Areg, Spp1, Inhba*), Endothelial cells (*Pecam1, Cdh5, Kdr, Cldn5, Sox17*), CAFs (*Col1a1, Fap, Pdgfra, Postn*), Pericytes (*Rgs5, Pdgfrb, Kcnj8, Des, Notch3*), NK cells (*Ncr1, Klrb1c, Eomes, Nkg7, Prf1*), gd T-cells (*Trdc, Trgc1, Il17a, Rorc*), Tregs (*Cd3e, Foxp3, Il2ra, Ctla4, Ikzf2, Tnfrsf18*), CTLs (*Cd3e, Cd8a, Cd8b1, Gzmb, Prf1*), Th cells (*Cd3e, Cd4, Il7r, Tcf7*), B cells (*Ccr7, Cd79a, Cd19, Ms4a1, Pax5, Ighd*), pDCs (*Siglech, Bst2, Tcf4, Spib, Irf7*). One cluster driven entirely by a single public accession was removed as sample specific. The annotated object was used both for downstream single-cell analyses and as the deconvolution reference.

### Slide-seq sample preparation and library generation

Epithelial and mixed tumors were dissected and embedded in optimal cutting temperature (OCT) compound and incubated on ice for 1 minute prior to submersion in an isopentane bath within a liquid nitrogen dewar. Frozen tumor tissue sections (10 μm thick) were mounted on 1 x 1 cm Spatial Transcriptomics Seeker tiles (Takara Bio) and sequencing libraries were generated according to the manufacturer’s instructions. Briefly, RNA capture and in situ reverse transcription were followed by enzymatic tissue digestion and release of the spatially barcoded beads. Second-strand cDNA synthesis and amplification were then performed, and sequencing libraries were prepared using the Nextera XT DNA kit. Libraries were pooled and sequenced on an Illumina NovaSeq X Plus instrument with 100 bp paired-end reads and 8 bp dual-index reads (Read 1/Read 2: 100 bp; Index 1/Index 2: 8 bp). Raw sequencing data were processed with the *slide_snake* pipeline (https://github.com/mckellardw/slide_snake). Adapter trimming and quality filtering were performed using cutadapt (v4.6)^98^ with the seeker_v3.1_internalTrim configuration. Reads were aligned to the mouse reference genome GRCm39 using STARsolo (v2.7.10b), with GENCODE M32 gene annotations, to generate gene-by-bead expression matrices^99^.

### Slide-seq data preprocessing

Per-sample spatial objects were concatenated on the union of genes and filtered to spots with more than 100 total counts. Sparse tissue edges and dissociated debris were then removed per sample: for each sample, a KD-tree was built on spot coordinates and the median distance to the 20th nearest neighbor was used as an estimate of local spot spacing. Spots with fewer than 30 neighbors within five times this median distance were discarded. One sample was exempted from this filter because its spot density profile differed from the others and the filter removed intact tissue.

### Spatial cell type deconvolution

Cell type composition was inferred at each spot with cell2location^100^. The raw annotated single-cell dataset was used as reference after mitochondrial, ribosomal and hemoglobin genes were removed. Reference genes were selected with the cell2location filtering utility (cell count cutoff of the larger of 20 cells or 0.1% of cells, cell percentage cutoff 0.005, non-zero mean cutoff 1.2). A negative binomial regression model was fit with the accession as batch covariate and the refined annotation as the cell type label, trained for 2,000 epochs with a batch size of 10,000, and the posterior was sampled 1,000 times to obtain per-cell-type reference signatures.

Genes were intersected for the spatial and reference to have a matching gene set. The spatial model was fit separately for each sample with the expected number of cells per location set to 1 and detection alpha set to 20, trained for 1,000 epochs with a batch size of 5,000 on the full data. Posterior 5% quantile abundances were exported (1,000 posterior samples) and row-normalized to per-spot cell type proportions. Each spot was additionally assigned a dominant label corresponding to its highest-proportion cell type, which was used for cell-type-resolved spatial analyses.

### Cellular neighborhood identification

Cellular neighborhoods were defined from local cell type composition. A per sample radius-neighbor graph was built on spot coordinates with a 50-unit radius and the spot itself included, and cell type proportions were summed across each spot’s neighborhood and renormalized. The neighborhood composition matrix was centered log-ratio transformed to account for its compositional nature, reduced by PCA to 14 components, and embedded with a 30-neighbor graph and UMAP. The final niche assignment was defined as the leiden clusters of 0.3 resolution. Niche composition was summarized as the mean cell type proportion per niche.

### Cell type co-localization

To determine cell type co-localization, pairwise spatial proximity between cell types was quantified by nearest-neighbor distance per sample, a KD-tree was built per cell type, and for every spot of cell type A the Euclidean distance to the nearest spot of cell type B was computed. To correct for differences in tissue size and spot density, distances were divided by the median nearest-neighbor distance among all spots in that sample. Then distances were aggregated as the median across all A spots to give a cell type by cell type distance matrix per sample. To focus on the quasi-mesenchymal compartment broadly, cycling and non-cycling tumor populations were merged, and the empirical cumulative distribution of distances from quasi-mesenchymal spots to TAMs, regulatory T cells, helper T cells, cytotoxic T cells and epithelial tumor cells was plotted, restricting to reference spots within 20 normalized distance units of a target and reporting the fraction of reference spots meeting that criterion.

### Gaussian smoothing of spatial expression

For use in downstream analysis, the spatial expression was denoised with Smoothie^60^. Spots with less than 50 total counts and genes with less than 100 counts tissue-wide were excluded. Then spots were normalized to 1,000 counts and log1p transformed. In-place Gaussian smoothing was applied to denoise the data at every spot with a kernel standard deviation of 40 coordinate units and a minimum of 25 spots within the kernel support.

Since smoothing over all spots mixes signal across adjacent cell types, a second smoothing pass was run on epithelial tumor spots alone, using the same kernel and a reduced minimum of 5 spots to accommodate the sparser support of a single compartment point set. This results in a clean non-contaminated epithelial expression surface. For distance-resolved analyses, the two representations were combined: rows of the whole-tissue smoothed matrix corresponding to epithelial tumor spots were replaced with their epithelium-only smoothed values, leaving all other spots unchanged.

### Epithelial cancer cell expression as a function of distance to quasi-mesenchymal cancer cells

To evaluate whether epithelial expression varies based on quasi-mesenchymal tumor cells proximal, spots were labeled as epithelial, quasi-mesenchymal or other using the dominant deconvolution label, and the merged epithelium-resolved smoothed matrix was used. For each epithelial spot, the Euclidean distance to the nearest quasi-mesenchymal spot was computed and normalized by the median nearest-neighbor distance among all spots in the sample. To prevent lone quasi cells from anchoring distances, quasi cells were only accepted as an anchor if at least three other quasi-mesenchymal spots lay within three normalized distance units. Epithelial spots more than 15 normalized units from any valid anchor were excluded.

Genes of high detection (in at least 5% of epithelial cells) were tested for association with distance by Spearman and Pearson correlation, with Benjamini-Hochberg correction across genes. Expression-distance relationships were visualized as LOWESS fits (smoothing fraction 0.35) of z-scored expression against normalized distance, with spots additionally binned by distance. Gene modules were evaluated the same way, scored either by control-matched gene set scoring (50 control genes, 25 expression bins) or by the first principal component of module genes computed within epithelial spots, with PC1 variance explained and gene loadings reported as a coherence check. Modules tested comprised MHC class I genes (*H2-K1*, *H2-D1*, *H2-Q7*) and MHC class I regulators (*Tap1*, *Tap2*, *Psmb8*, *Psmb9*, *Nlrc5*).

### RNA sequencing of FACS-isolated epithelial cells

At least 10,000 CD45^neg^ Snail-YFP^neg^ EpCAM^hi^ epithelial cancer cells were sorted from pB2 and pB2:Snail^hi^ 9:1 mixed tumors following the gating strategy described in Fig S6A and using a BD Biosciences FACSMelody Cell Sorter. Sorted cells were pelleted by centrifugation at 1250 rpm for 5 minutes at 4°C. RNA was extracted using the Norgen Biotek Total RNA Purification Kit, followed by DNAse treatment. RNA sample quality was confirmed using a Qubit (RNA HS kit; Thermo Fisher) to determine concentration and with Femto Pulse (Agilent) to determine RNA integrity. Total RNA samples were treated with a FastSelect rRNA depletion kit (Qiagen) to remove rRNA. UDI-barcoded RNAseq libraries were generated with the NEBNext Ultra II Nondirectional RNA Library Prep Kit (New England Biolabs). Each library was quantified with a Qubit (dsDNA HS kit; Thermo Fisher) and the size distribution was determined with a Fragment Analyzer (Agilent) prior to pooling. Libraries were sequenced on an Illumina instrument. At least 20M reads were generated per library.

Reads were trimmed for low quality and adapter sequences with fastp v0.21.0^101^. Reads were mapped to the reference genome/transcriptome (Ensembl GRCm38) using STAR v2.7.0e^102^. SARTools and DESeq2 v1.26.0 were used to generate normalized counts and statistical analysis of differential gene expression^103,104^.

### TNBC spatial data processing and normalization

Spatial transcriptomics data from multiple patient samples were imported and merged into a unified dataset using Scanpy. Quality control was performed by filtering out cells with fewer than 500 total transcript counts. Gene expression data were then normalized to 10,000 counts per cell and log1p-transformed for downstream analysis.

### Pathway activity scoring

Gene signature activities, including Epithelial-Mesenchymal Transition (EMT), were obtained from MSigDB^69^ and quantified at single-cell resolution. The AUCell^105^ algorithm, implemented via the decoupler^106^ package, was utilized to compute pathway enrichment scores. Mean pathway activity scores were subsequently aggregated for the tumor cell population within each patient sample.

### Spatial proximity and infiltration analysis

To evaluate the spatial distribution of the immune infiltrate relative to the tumor parenchyma, k-d tree spatial indexing^107^ (SciPy) was applied to physical cellular coordinates. For each immune cell state (encompassing T cell and myeloid lineages), two spatial metrics were calculated per patient: 1) the overall median distance from the target immune cells to the nearest tumor cell and 2) an infiltration ratio, defined as the proportion of target immune cells located within a 30 µm radius of the nearest tumor cell.

### Statistical analysis

All statistical analyses were performed using the GraphPad Prism v10.6.0 software, apart from analysis of human TNBC spatial data. Bar plots and tumor kinetics graphs indicate mean value with error bars representing standard error of the mean (SEM). Count data were analyzed using two-tailed unpaired t-tests (Fig 1B-F, 1K-L, 3A, 3E-F, 4C, 5A, S1A-F, S2B-J, S3A-L, S6B, S7G, S8A, S8E, S8G, S9C), one-tailed Mann-Whitney test (Fig 4D-E, S8C, S9D, S9F) one-way ANOVA (Fig 2A-B, 4C, 5D, S7D, S7G, S8D), or a Kruskal-Wallis test (Fig 4F, 5E-F, S8D, S9A, S9H). Tumor kinetics were analyzed with mixed-effects model (Fig 1K-L, 2A-B, 3E, 5E, S1A, S1F, S2J, S7F, S7G, S8E, S8G). Asterisks indicate statistical significance where \**p* < 0.05, \*\**p* < 0.01, \*\*\**p* < 0.001, \*\*\*\**p* < 0.0001. Comparisons are not significant (*p* ≥ 0.05) if not indicated. To identify associations between EMP activation and immune cell spatial distribution in human TNBC samples, Pearson and Spearman correlation coefficients were calculated between patient-level mean tumor pathway scores and immune cell spatial metrics. P-values were adjusted for multiple hypothesis testing using the Benjamini-Hochberg false discovery rate (FDR) procedure. Correlations with an FDR < 0.05 were considered statistically significant (*).

## Data Availability

All raw sequencing data, spatial barcodes lists, and raw count matrices will be made publicly available upon publication.

## Code Availability

All scripts and code used to analyze the scRNA-seq and spatial data will be made publicly available on GitHub upon publication.

## Acknowledgments

We are grateful to all members of the Dongre Lab for their help in discussing and reviewing the manuscript. We thank the Center for Animal Resources and Education (CARE) at Cornell University for taking care of the mice used in these studies. We are grateful to Dr. Yassi Hafezi of the Cornell University Biotechnology Resource Center Genomics Facility (RRID: SCR_021727) for generating the scRNA-seq data. We thank Jen Grenier of the Cornell Transcriptional Regulation and Expression (TREx) facility (RRID: SCR_022532) for her help in generating the RNA sequencing data. Additionally, we thank staff at the Cornell Institute of Biotechnology Flow Cytometry Facility (RRID: SCR_021740) for their assistance in generating flow cytometry data presented in this study. We thank staff in the Cornell University Animal Health Diagnostic Laboratory for processing tissue samples for histology. Funding for this study was provided by the K22 Transition Career Development Award (A.D.), Judith Appleton, Ph.D., Early Career Excellence in Research Award (A.D.), Cornell Center for Immunology seed grant (A.D. and I.D.V.), and Cornell University start-up funds (A.D.). M.K.J and S.S were supported by Param Hansa Philanthropies.

## Author Contributions

Conceptualization: KB, AD

Methodology: KB, AS, SS, IN, RP, SB, PS

Investigation: KB, AS, SS, IN, KP, MR, RP, SB, PS, SC

Visualization: KB, AS, SS

Supervision: MKJ, IDV, PS, AD

Writing—original draft: KB

Writing—review & editing: KB, AD

