## Supplemental Figures for "MHC-I downregulation is associated with immune-evasion in breast cancers with heterogeneous activation of epithelial-mesenchymal plasticity"

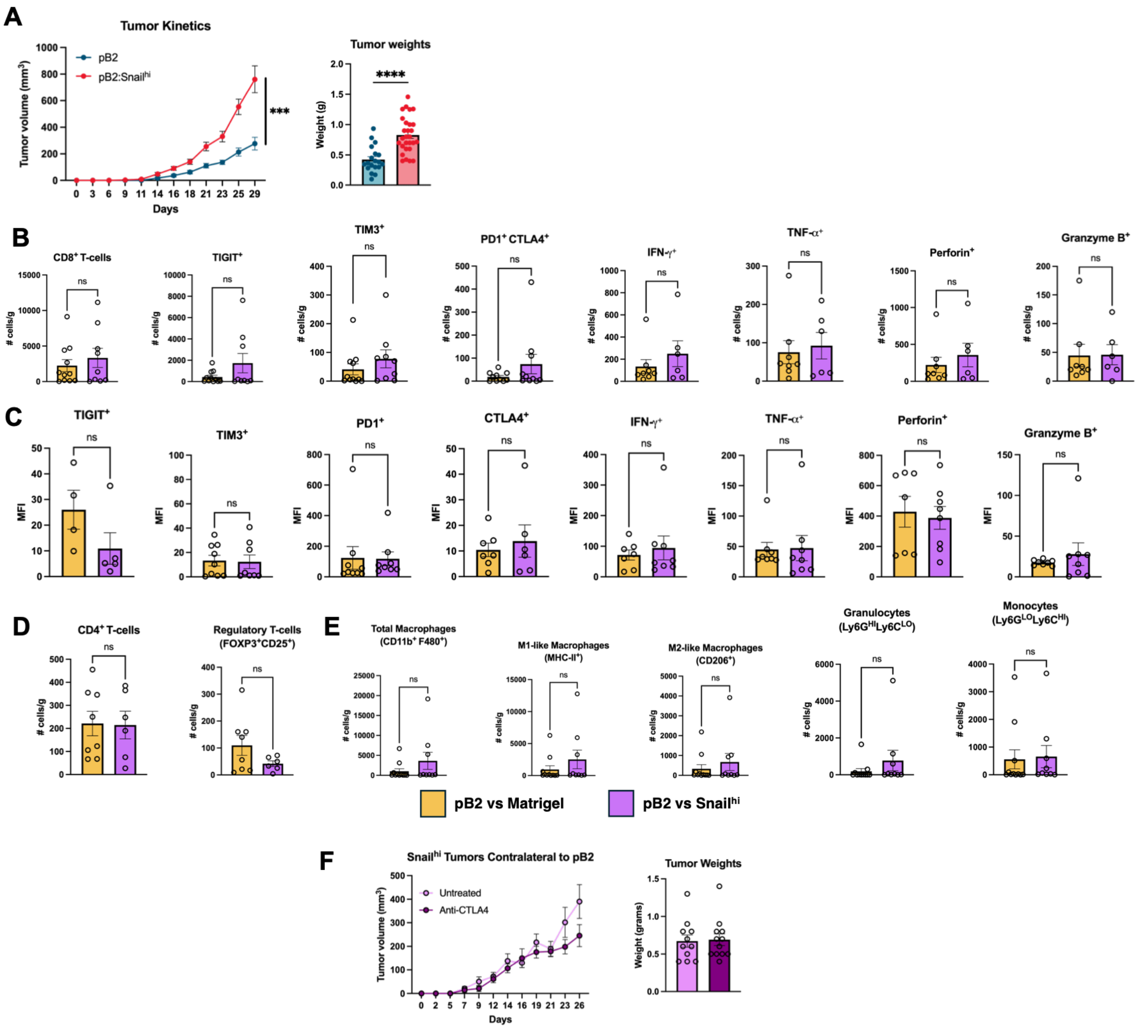


**Figure S1. Additional flow cytometry and tumor kinetic data from pB2 contralateral model. (A)** Tumor kinetics and weights of untreated pB2 and pB2:Snail$^{hi}$ mixed tumors. **(B)** Number of cells/gram of tumor weight (# cells/g) and **(C)** median fluorescent intensity (MFI) data for flow cytometry data presented in Figure 1 quantifying CD8$^{+}$ T-cell markers in pB2 tumors contralateral to Matrigel (Mg, yellow) or Snail$^{hi}$ tumor (purple). **(D)** Cells/g for Tregs, and **(E)** myeloid cells in pB2 tumors contralateral to Mg (yellow) or Snail$^{hi}$ tumor (purple). **(F)** Tumor kinetics and weights for Snail$^{hi}$ tumors implanted contralateral to pB2 tumors in Fig 1L. **A** represents six independent experimental replicates with *n*=3-5 per group. **B-E** include three independent experimental replicates including *n*=3-4 per group. **F** includes three independent experimental replicates including *n*=3-4 per group. Data are represented as mean ± standard error of the mean (SEM) and were analyzed using an unpaired two-tailed t-test (**A-F**) or mixed-effects model (tumor kinetics in **A** and **F**). ***$p < 0.001$, ****$p < 0.0001$. Comparisons are not significant (ns, $p \geq 0.05$) if not indicated.

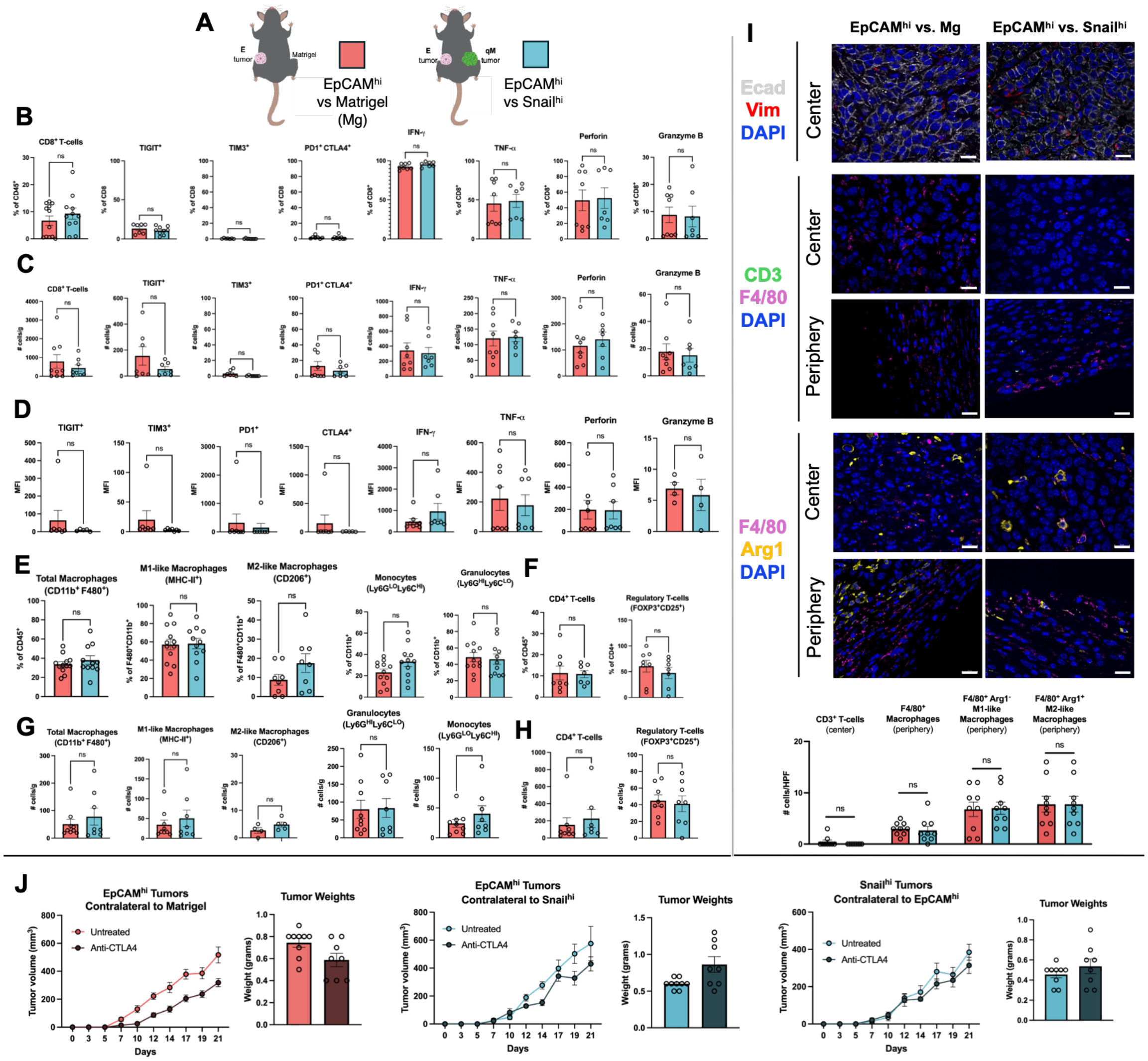


**Figure S2. Contralateral model using EpCAM^hi^ cells. (A)** EpCAM^hi^ cells were implanted contralateral to Matrigel (Mg) or Snail^hi^ cells. **(B)** Percentages, **(C)** # of cells/g, and **(D)** median fluorescent intensity (MFI) of $CD8^+$ T-cell markers within EpCAM^hi^ tumors contralateral to Mg or Snail^hi^ cells, as measured by flow cytometry. **(E-F)** Percentages and **(G-H)** # of cells/g of myeloid and $CD4^+$ T-cells in EpCAM^hi^ tumors. **(I)** Immunofluorescent staining of EpCAM^hi^ tumors contralateral to Mg or Snail^hi^ for EMP markers (E-cadherin, Ecad, white; Vimentin, Vim, red) and immune cell markers (CD3- T-cells, green; F4/80- macrophages, magenta; Arginase-1, Arg1, yellow). Scale bars indicate 20 μm. **(J)** Tumor kinetics and weights for EpCAM^hi^ tumors contralateral to Mg or Snail^hi^ tumors and treated with anti-CTLA-4. Data in **A-H** represent three independent experimental replicates with *n*=4 per group. **I** includes two experimental replicates with total *n*=3 per group. **J** represents two independent experimental replicates with *n*=3-5 per group. Data are represented as mean ± standard error of the mean (SEM) and were analyzed using an unpaired two-tailed t-test (**A-J**) or mixed-effects model (tumor kinetics in **J**). Comparisons are not significant (ns, $p \geq 0.05$) if not indicated.

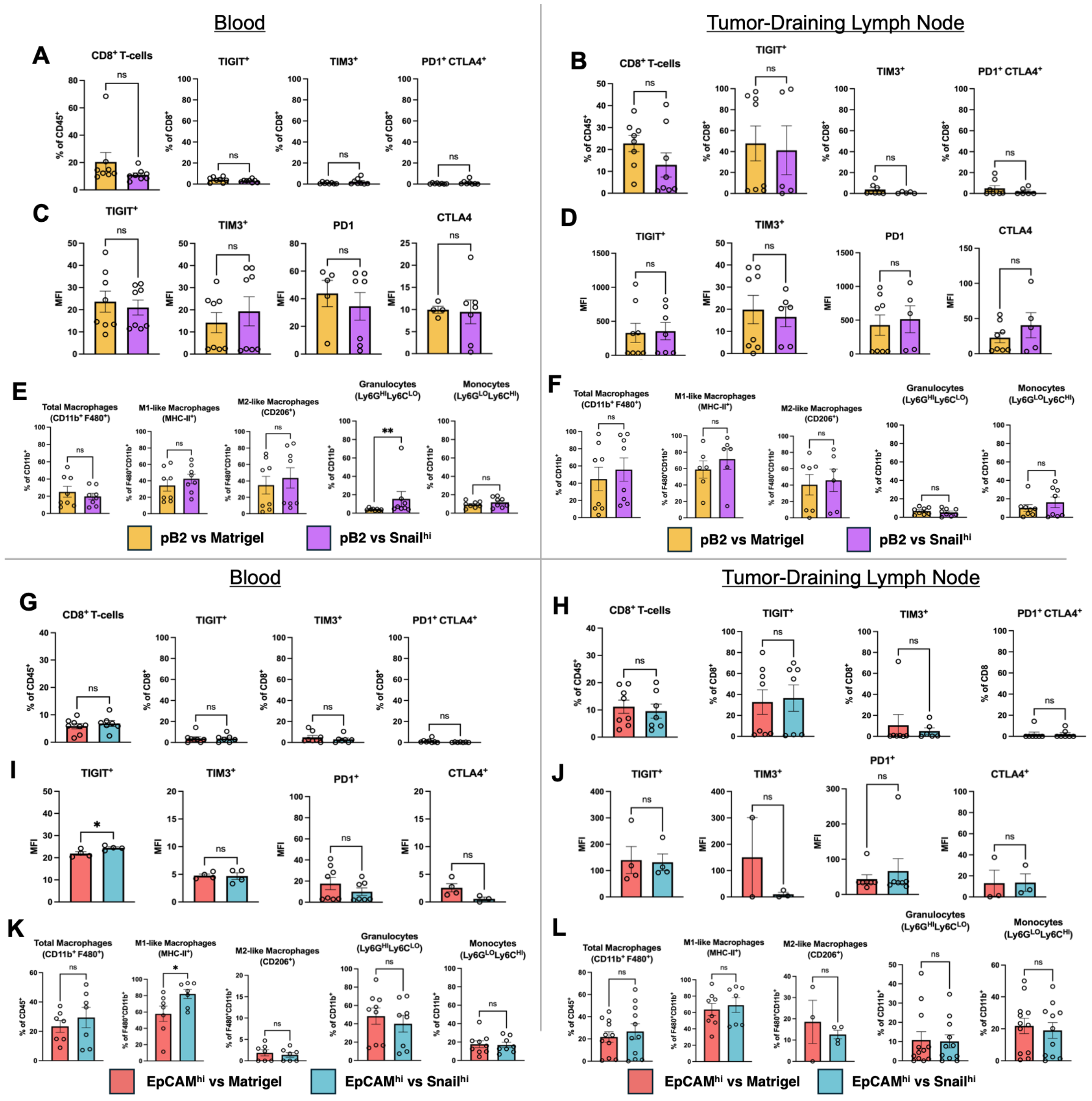


**Figure S3. Immune cell markers in blood and tumor-draining lymph nodes from contralateral models. (A-B)** Percentage and **(C-D)** median fluorescent intensity (MFI) of CD8$^{+}$ T-cell dysfunction markers in the blood and tumor-draining lymph node of mice bearing pB2 tumors contralateral to Matrigel (Mg) or Snail$^{hi}$ tumor. **(E-F)** Myeloid cell markers in the blood and tumor-draining lymph node of mice bearing pB2 tumors contralateral to Mg or Snail$^{hi}$ tumor**. (G-H)** Percentage and **(I-J)** MFI of CD8$^{+}$ T-cell dysfunction markers in the blood and tumor-draining lymph node of mice bearing EpCAM$^{hi}$ tumors contralateral to Mg or Snail$^{hi}$ tumor. **(K-L)** Myeloid cell markers in the blood and tumor-draining lymph node of mice bearing EpCAM$^{hi}$ tumors contralateral to Mg or Snail$^{hi}$ tumor. Data in **A-L** represent two independent experimental replicates with $n$=4 per group. Data are represented as mean ± standard error of the mean (SEM) and were analyzed using an unpaired two-tailed t-test. **$p < 0.01$. Comparisons are not significant (ns, $p \geq 0.05$) if not indicated.

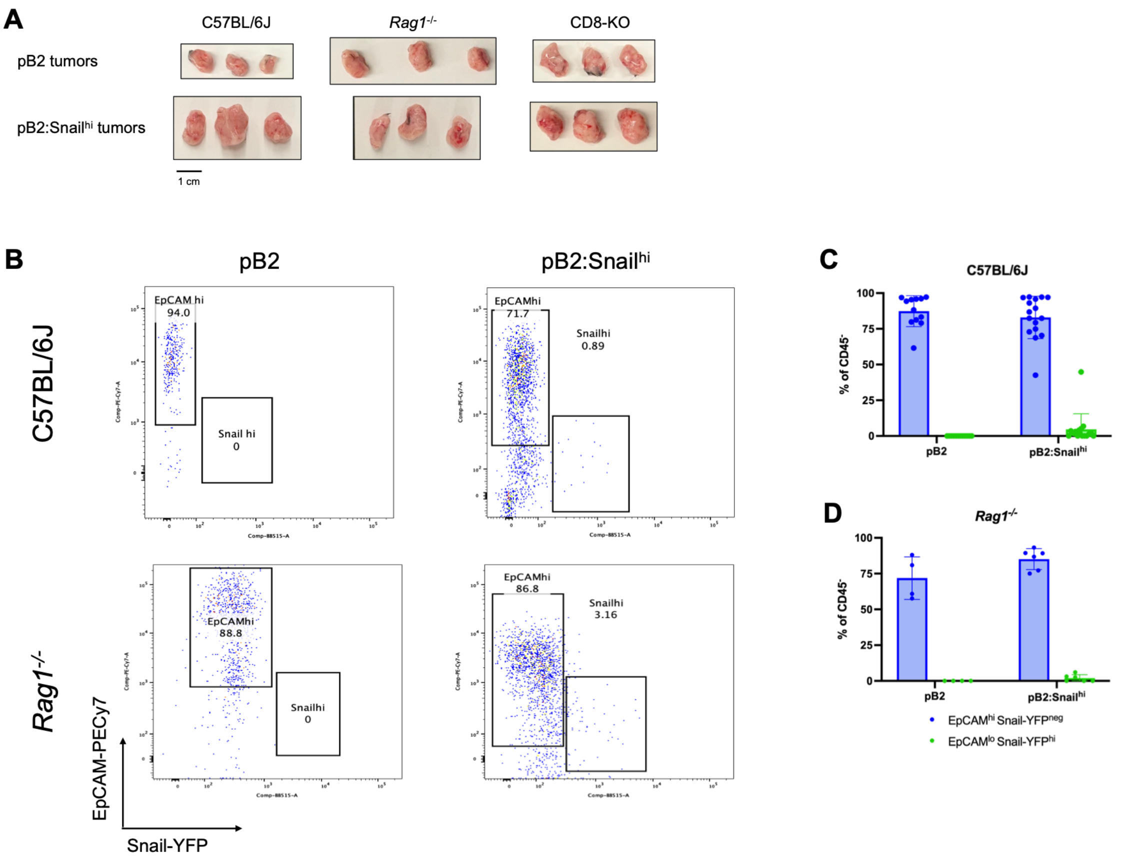


**Figure S4. Additional data from mixed tumors implanted in immunodeficient hosts.** **(A)** Representative images of tumors grown in C57BL/6J, *Rag1*$^{-/-}$, or CD8-KO mice. **(B)** Representative flow cytometry plots of EpCAM$^{hi}$ Snail-YFP$^{neg}$ epithelial cancer cells and EpCAM$^{lo}$ Snail-YFP$^{hi}$ quasi-mesenchymal cancer cells in pB2 and pB2:Snail$^{hi}$ tumors. **(C-D)** Quantification of percentage of epithelial and quasi-mesenchymal cancer cells in pB2 and pB2:Snail$^{hi}$ tumors implanted in C57BL/6J or *Rag1*$^{-/-}$ mice. Data in **C** represent four experimental replicates with *n*=3-4 per group. **D** includes one or two experimental replicates with *n*=3-4 per group. Data are represented as mean ± standard error of the mean (SEM).

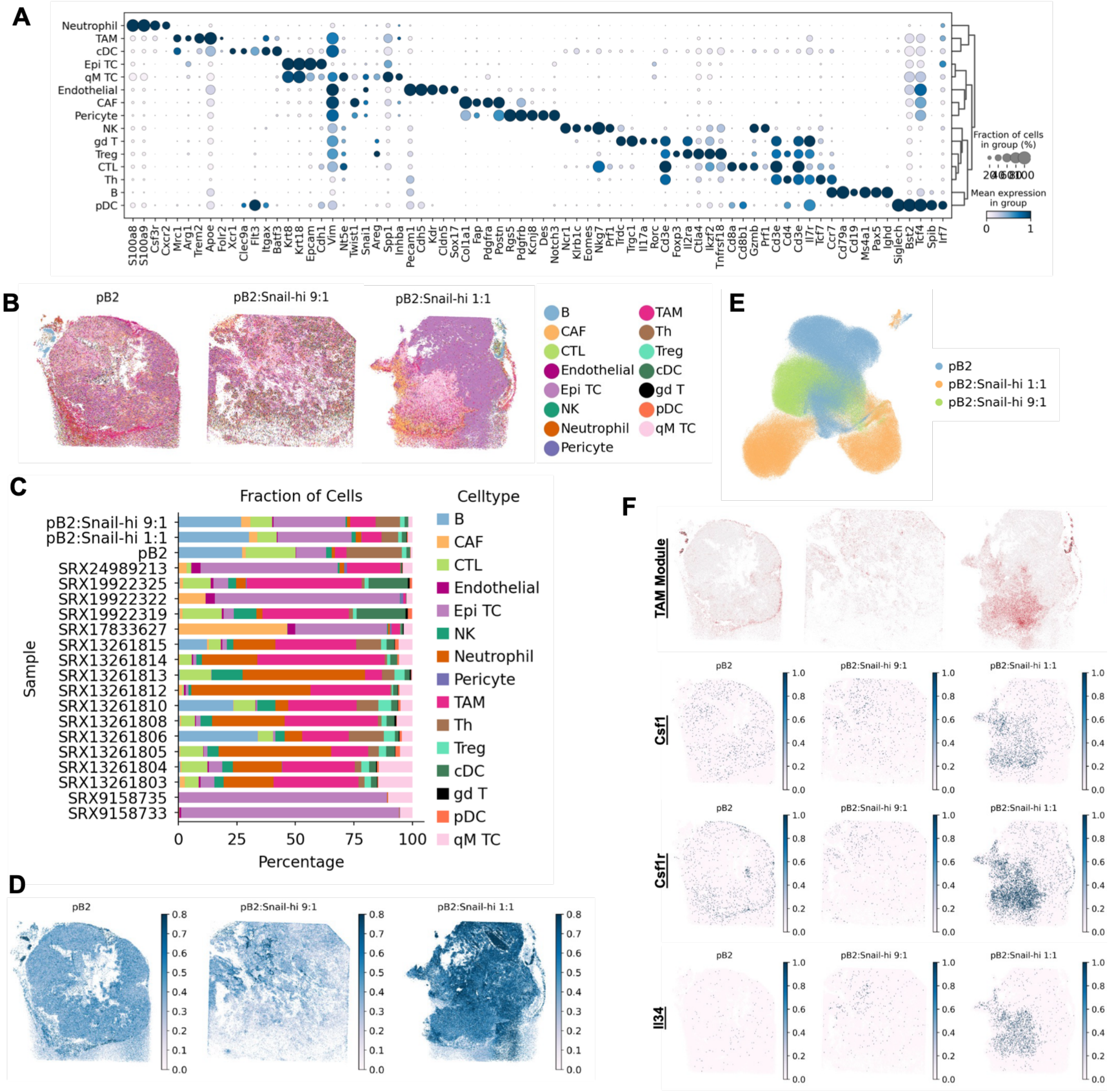


**Figure S5. Single-cell RNA sequencing and spatial transcriptomics of epithelial and mixed tumors. (A)** Marker genes used to annotate single-cell RNA sequencing (scRNA-seq) clusters. **(B)** Cell2location cell type prediction of Slide-seq samples using the scRNA-seq data as reference. **(C)** Proportions of cell types in scRNA-seq of epithelial and mixed tumors split by sample. **(D)** Maximum prediction score from Cell2location analysis. **(E)** Sample makeup of cellular neighborhoods. **(F)** Tumor-associated macrophage (TAM) module identified using Smoothie analysis and expression of *Csf1*, *Csf1r*, and *Il34* in spatial transcriptomics data. All data represent one replicate per condition.

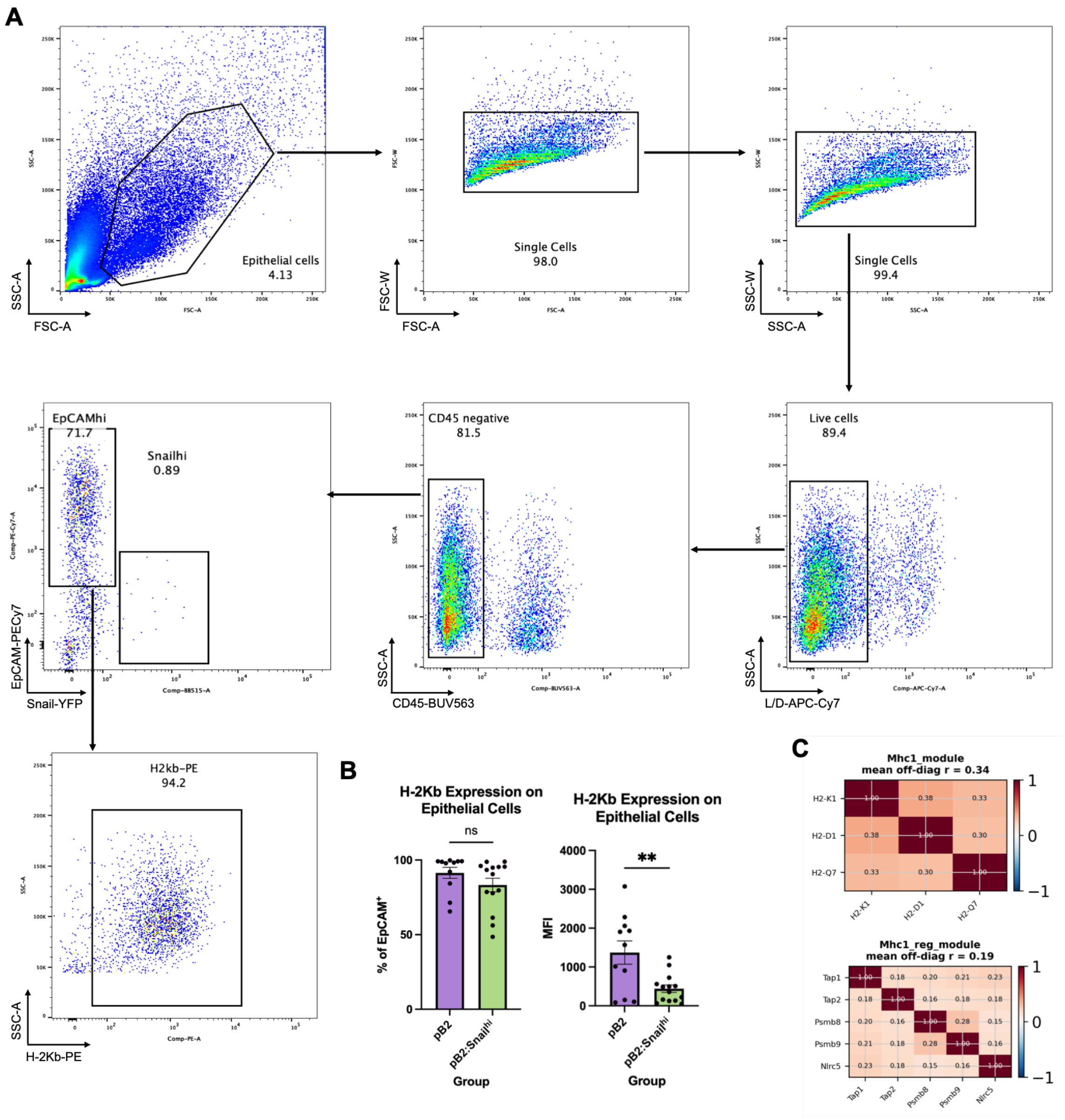


**Figure S6. Approach for measuring H-2Kb on epithelial cancer cells by flow cytometry. (A)** Visualization of gating strategy used to analyze flow cytometry data from mixed tumors to quantify H-2Kb expression on epithelial cells within mixed tumors. **(B)** Percentage of epithelial cancer cells expressing H-2Kb and median fluorescent intensity in pB2 and pB2:Snail$^{hi}$ tumors. **(C)** Pairwise Spearman correlation between MHC class I (top) and antigen presentation (bottom) module genes across all Epi spots. Mean off-diagonal r summarizes module coherence. **B** represents four independent experimental replicates including *n*=3-5 per group. Data are represented as mean $\pm$ standard error of the mean (SEM) and were analyzed using an unpaired two-tailed t-test. $^{**}p < 0.01$, ns $p \geq 0.05$.

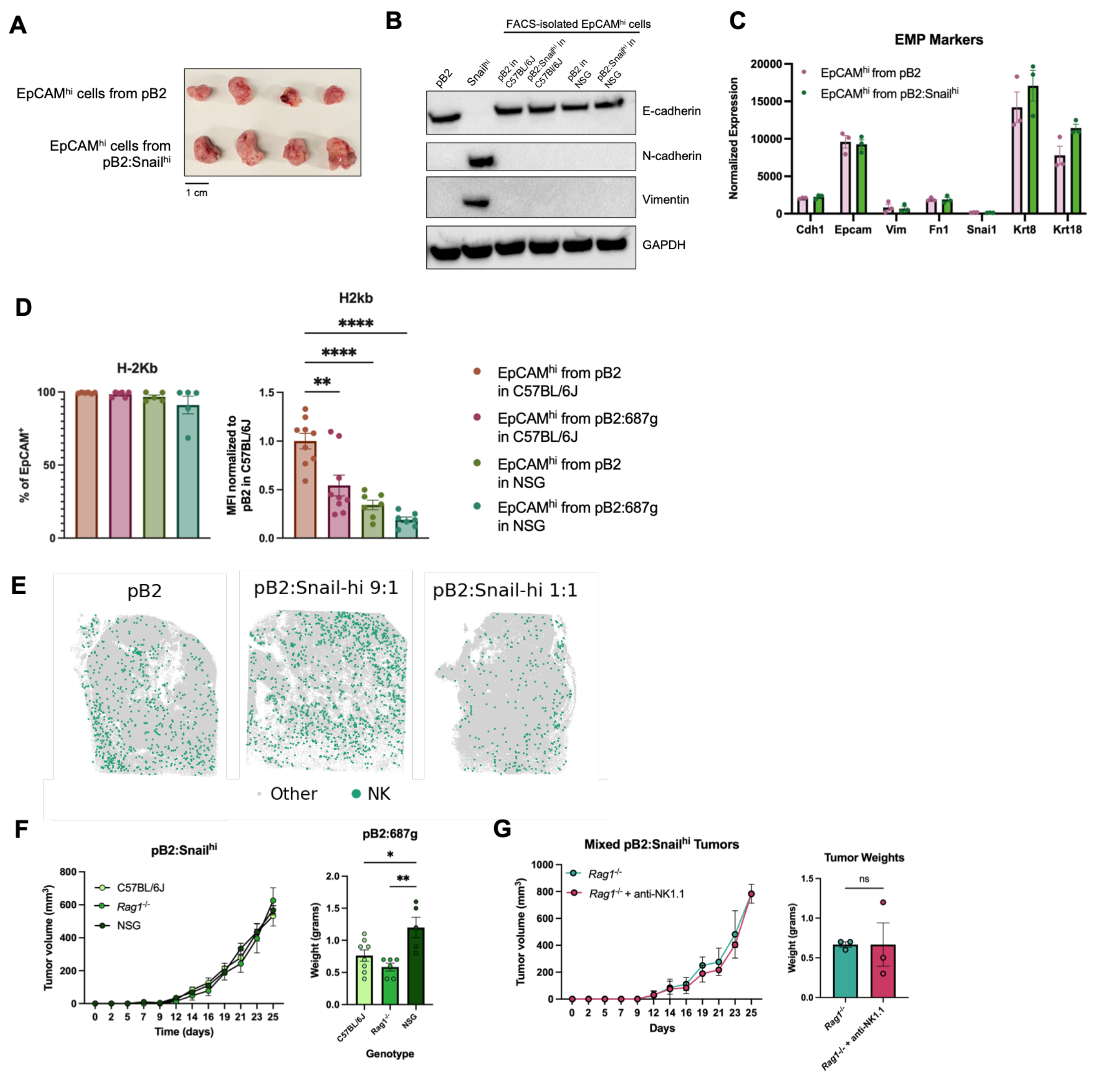


**Figure S7. Additional data from epithelial reimplantation experiments and role of NK cells in mixed tumors. (A)** Representative images of tumors resulting from reimplantation of epithelial cancer cells isolated from pB2 or pB2:Snail$^{hi}$ tumors grown in C57BL/6J mice. **(B)** Western blot demonstrating that FACS-isolated epithelial cancer cells remain epithelial prior to reimplantation. **(C)** Normalized expression of EMP markers in epithelial cells isolated from pB2 or pB2:Snail$^{hi}$ tumors. **(D)** Percentage of epithelial cancer cells expressing H-2Kb and median fluorescent intensity (MFI) from epithelial cells isolated from pB2 or pB2:Snail$^{hi}$ tumors grown in C57BL/6J or NSG mice and reimplanted in C57BL/6J mice. **(E)** Cell2location predicted location of NK cells in Slide-seq samples. **(F)**

Tumor kinetics and weights of pB2:Snail$^{hi}$ tumors implanted in C57BL/6J, *Rag1*$^{-/-}$, and NSG mice. **(G)** Tumor kinetics and weights from NK cell-depleted mixed tumor-bearing *Rag1*$^{-/-}$ mice. **C** represents one experimental replicate with *n*=3 per group. **D** includes two or three independent experimental replicates with *n*=3-4 per group. **F** represents two experimental replicates with *n*=3-4 per group. **G** includes one experimental replicate with *n*=3 per group. Data in **C, D, F,** and **G** are represented as mean $\pm$ standard error of the mean (SEM). Data in **D** and **F** (tumor weights, right) were analyzed using a one-way ANOVA. Tumor kinetics data in **F** and **G** were analyzed using a mixed-effects model. An unpaired two-tailed t-test was used to compare tumor weights in **G.** *$p < 0.05$, **$p < 0.01$, ****$p < 0.0001$. Comparisons are not significant (ns, $p \geq 0.05$) if not indicated.

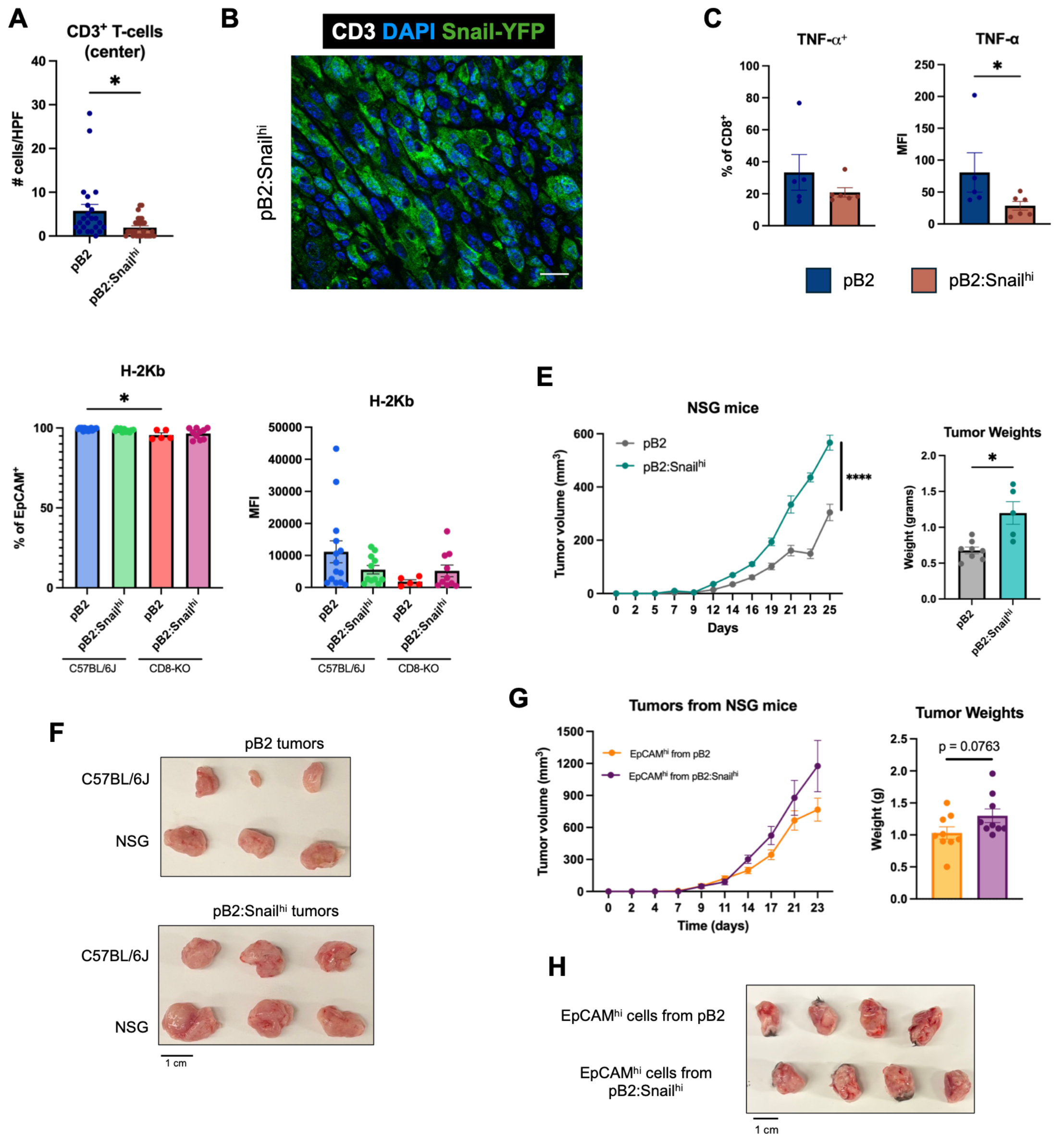


**Figure S8. Role of CD8$^{+}$ T-cells in mixed tumors and implantation in NSG mice. (A)** Quantification of total intratumoral T-cells in pB2 and pB2:Snail$^{hi}$ tumors. **(B)** Representative image of T-cells (white) in Snail$^{hi}$ region (green) of a pB2:Snail$^{hi}$ tumor. Scale bar indicates 20 μm. **(C)** Expression of TNF-α in pB2 and pB2:Snail$^{hi}$ tumors. **(D)** Percentage of H-2Kb-expressing epithelial cancer cells and absolute MFI in pB2 and pB2:Snail$^{hi}$ tumors implanted in C57BL/6J or CD8-KO mice. **(E)** Tumor kinetics and weights of pB2 and pB2:Snail$^{hi}$ tumors grown in NSG mice. **(F)** Representative images of pB2 and pB2:Snail$^{hi}$ tumors grown in C57BL/6J or NSG mice. **(G)** Tumor kinetics and weights from reimplantation of epithelial cancer cells isolated from pB2 or pB2:Snail$^{hi}$ tumors grown in NSG mice. **(H)** Representative images of tumors formed from reimplantation of epithelial cancer cells isolated from pB2 and pB2:Snail$^{hi}$ tumors grown in NSG mice. **A-B** represent two independent experimental replicates including $n$=5 total per group. **C** includes two independent experimental replicates with $n$=2-3 per group. **D** includes two or three independent experimental replicates with $n$=3-5 per group. **E** includes two experimental replicates with $n$=3-4 per group. **G** includes three experimental replicates with $n$=3 per group. Data in **A, C, D, E,** and **G** are represented as mean $\pm$ standard error of the mean (SEM). Data in **A**, **E,** and **G** were analyzed using an unpaired two-tailed t-test, Data in **C** were analyzed using a one-tailed Mann-Whitney test. Data in **D** were analyzed with a Kruskal-Wallis test. Tumor kinetics data in **E** and **G** were analyzed using a mixed-effects model. *$p < 0.05$. Comparisons are not significant (ns, $p \geq 0.05$) if not indicated.

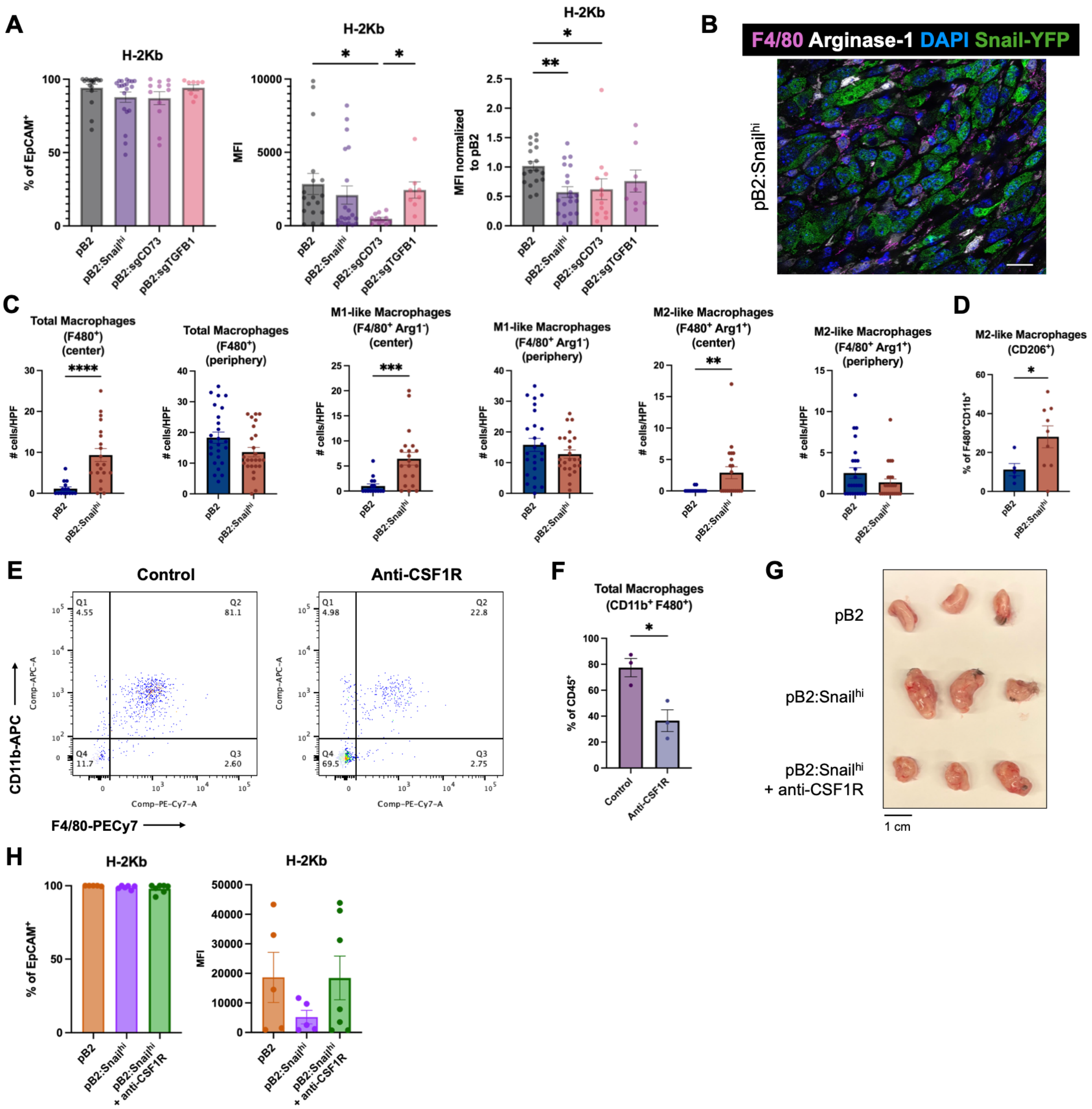


**Figure S9. T-cell independent MHC-I downregulation on E cancer cells. (A)** Percentage of epithelial cancer cells expressing H-2Kb and median fluorescent intensity in mixed tumors containing CD73 or TGF-$\beta$1-deficient quasi-mesenchymal cancer cells. **(B)** Representative image of Arginase-1$^+$ (white) F4/80$^+$ (magenta) macrophages in Snail$^{hi}$ region (green) of a pB2:Snail$^{hi}$ tumor. Scale bar indicates 20 $\mu$m. **(C)** Additional macrophage quantification data from immunofluorescent staining of pB2 and pB2:Snail$^{hi}$ tumors. **(D)** Percentage of CD206+ M2-like macrophages in pB2 and pB2:Snail$^{hi}$ tumors, as measured by flow cytometry. **(E)** FACS plots demonstrating macrophage depletion upon anti-CSF1R treatment in the primary tumor and spleen. **(F)** Quantification of macrophage depletion. **(G)** Representative images of mixed pB2:Snail$^{hi}$ tumors with or without macrophage depletion in comparison to epithelial pB2 tumors. **(H)** Percentage of H-2Kb-expressing cancer cells in macrophage-depleted mixed tumors. **A** represents three (pB2:sgTGFB1) or four (pB2:sgCD73) independent experimental replicates including $n$=3-5 per group. **B** represents two independent experimental replicates including $n$=5 total per group. **E-F** represent one experimental replicate with $n$=3 per group. **G-H** represents two experimental replicates with $n$=2-3 per group. Data in **A, C, D, F,** and **H** are represented as mean $\pm$ standard error of the mean (SEM). Data in **A** were analyzed using a Kruskal-Wallis test. Data in **C** were analyzed using an unpaired two-tailed t-test. Data in **D** were analyzed using a one-tailed Mann-Whitney test. Data in **F** were analyzed with a Mann-Whitney test. Data in **H** were analyzed using a Kruskal-Wallis test. *$p < 0.05$, **$p < 0.01$, ***$p < 0.001$, ****$p < 0.0001$. Comparisons are not significant (ns, $p \geq 0.05$) if not indicated.

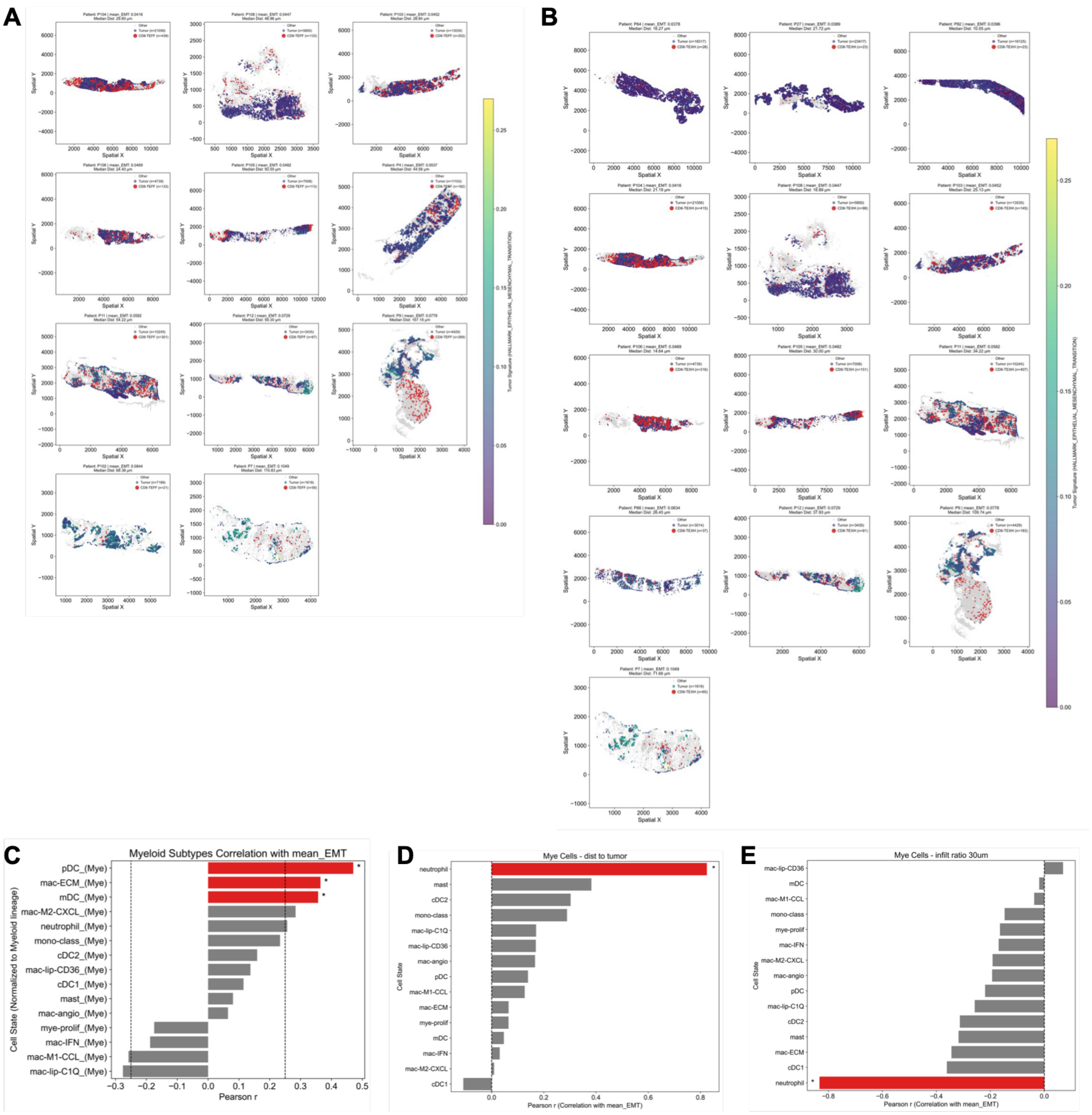


**Figure S10. Additional data from analysis of human triple-negative breast cancer spatial samples. (A)** Distribution of effector CD8$^+$ T-cells in TNBC samples with increasing EMP activation. Tumor cells are colored by EMP enrichment and effector CD8$^+$ T-cells are depicted in red. **(B)** Distribution of exhausted CD8$^+$ T-cells in TNBC samples with increasing EMP activation. Tumor cells are colored by EMP enrichment and exhausted CD8$^+$ T-cells are depicted in red. **(C)** Correlation of the proportion of each myeloid cell subtype with mean epithelial-mesenchymal transition (EMT) score in tumor cells. **(D)** Correlation of the distance of each myeloid cell subtype from nearest tumor cell with mean epithelial-mesenchymal transition (EMT) score in tumor cells. **(E)** Correlation of the infiltration ratio of each myeloid cell subtype with mean epithelial-mesenchymal transition (EMT) score in tumor cells. **C-E** include 44 TNBC patient samples. In **C-E**, red bars and * indicate false discovery rate (FDR) < 0.05.
